# Label-free Isolation and Single Cell Biophysical Phenotyping Analysis of Primary Cardiomyocytes Using Inertial Microfluidics

**DOI:** 10.1101/2020.08.31.243592

**Authors:** Hossein Tavassoli, Prunella Rorimpandey, Young Chan Kang, Michael Carnell, Chris Brownlee, John E Pimanda, Peggy P.Y. Chan, Vashe Chandrakanthan

## Abstract

To advance our understanding of cardiomyocyte identity and function, we need appropriate tools to isolate pure primary cardiomyocytes. We have developed a label-free method to purify viable cardiomyocytes from mouse neonatal hearts using a simple inertial microfluidics biochip. Cardiomyocytes were sorted from neonatal hearts and isolated to >90% purity and their physico-mechanical properties were evaluated using real time deformability cytometry. Purified cardiomyocytes were viable and retained their identity and function as depicted by expression of cardiac specific markers and contractility. Furthermore, we showed that cardiomyocytes have a distinct physico-mechanical phenotype that could be used as an intrinsic biophysical marker to distinguish these cells from other cell types within the heart. Taken together, this cardiomyocyte isolation and phenotyping method could serve as a valuable tool to progress our understanding of cardiomyocyte identity and function, which will ultimately benefit many diagnostic development and cardiac treatment studies.

## 1. Introduction

Cardiovascular disease is the leading cause of death worldwide, outstripping death due to all cancers combined. The prevalence of end-stage heart failure is increasing [1, 2] and although heart transplantation is an option for patients when pharmacological therapies fail, donor hearts are scarce and alternatives that promote tissue regeneration are urgently needed. In this regard, neonatal mouse cardiomyocytes (CMs) are of interest because of their unique proliferative nature. A significant number of studies have shown that the mouse neonatal heart possesses regenerative characteristics [3, 4]. The physico-mechanical properties of CMs are not well characterised but benchmarking these properties are important for the development of tissue engineered cardiac biomaterials [5] and cell therapy cardiac regeneration [6].

Prior to studying physico-mechanical properties of neonatal CMs, these cells need to be extracted from the intact heart. Various methods have been used to isolate CMs, however the efficiency, purity and post-isolation functionality is highly variable [7]. One of the first described methods used to isolate CMs was pre-plating of collagenase-digested cardiac cell populations based on different rates of cell attachment [8, 9]. Although this was a simple method, drawbacks such as long processing time, poor cell type specificity, change in gene expression profiles and lack of consistency have made this method unreliable [10–12]. Other methods include cell isolation based on cell-surface antigens (such as VCAM1 [13], SIRPA [14], SSEA-1 [15] and ALCAM [16]) and intracellular markers (such as nanoscale probes called molecular beacons (MB), which target CM-mRNAs [17–20] or a fluorescent dye that labels CM mitochondria known as tetramethylrhodamine methyl ester perchlorate (TMRM) [21]). Labelled cells are sorted by fluorescence activated cell sorting (FACS) or magnetic activated cell sorting (MACS). Another isolation method relied on labelling CMs based on their chemical properties, where cardiac cells were treated with a sodium nitrite solution, making them paramagnetic prior to passing them through a magnetic field [22]. Recently, a similar method using treatment of cardiac cells with super-paramagnetic iron oxide particles has been described [12]. However, application of labelling techniques has limitations due to the requirement of expensive antibodies and auxiliary equipment, and potential alterations of cell identity. Furthermore, many of these published labelling methods are limited to CMs derived from induced pluripotent stem cells (iPSCs) [14, 15, 17, 21, 23–25] or embryonic stem cells (ESCs) derived CMs [14–18, 24, 26–29]. These cultured cells are distinct from primary cardiomyocytes with respect to their electrophysiological characteristics, structure, size, phenotype, contractility, membrane potential, and residual epigenetic memory and are not directly comparable [30]. Given that primary neonatal CMs are an essential tool for understanding cardiac development and regeneration, a label-free method that can isolate primary neonatal CM from an intact heart is highly desirable.

Microfluidic devices have emerged as attractive tools for cell separation and sorting [31]. Two microfluidic systems that demonstrated the feasibility of CM isolation and enrichment have been described [10, 11]. Murthy et al., [10] have demonstrated the use of a diffusive filtering microfluidic device that contained a main middle channel connected to two side channels with two micro-sieve structures (5 μm tall, 40 μm length) interfacing the channels. A mixture of non-myocytes and myocytes were isolated from ventricles of neonatal Sprague Dawley rat and then fed into the main channel at a flow rate of 20 μl/min. The micro-sieve structure functioned as a filter to trap larger myocytes in the middle channel and allowed smaller non-myocyte to flow into the side channel to obtain non-myocyte separation by sizes [10]. However, this method was susceptible to device clogging [11]. Zhang et al., [11] have demonstrated the use of a deterministic lateral displacement microfluidic device containing an array of posts and seven outlets. Neonatal CM from quartered hearts of Sprague-Dawley rats were first pre-filtered using a microfluidic filter to remove clogs. The cells were next fed to the array device at an inlet flow rate of 80 μl/min, smaller cells follow the fluid flow direction while larger cells follow the direction of the posts separated from the main flow by a small angle. Fractions of total cell output were collected at different outlet where size dependent separation could be found between two of the outlets generating a yield of around 55% [11].

Although both diffusive filtering devices and deterministic lateral displacement devices can separate cells passively without the use of label, the geometries of the micro-sieve and the array of posts need to be tailored differently for separation of cells with different size range [32–34]. Changing the geometries of these devices will require fabrication of new master moulds. However, fabrication of these master moulds requires high precision facilities and clean rooms, which limits their application to highly specialised laboratories. Furthermore, these devices have limited practical applications due to slow throughput flow rate. Hence, there is a need for a label-free, fast, higher throughput and cost-effective method to isolate functional primary CMs directly from the heart with minimal manipulation.

Inertial microfluidic devices with curved microchannels are able to concentrate dispersed particles into a narrow band. This focusing phenomenon is a result of a balance between shear gradient lift force and the wall effect lift force. Inertial cell separation is achieved by manipulating the differences in lift forces that are depending on particle size. One of the most attractive features of inertial microfluidic devices is that they are built with open channels with no blocking structures and utilise high liquid shear, thus minimise clogging tendency [31]. In addition, inertial devices can be operated with much higher flow rate (millilitre level) compared to diffusive filtering and lateral displacement devices, they are therefore very attractive for biological applications [35]. Inertial microfluidics has already been employed [36, 37] for various types of cell separation and cell sorting studies, such as filtration [38], blood fractionation [39, 40], bacteria detection [41], viral recovery [42], microalgae separation [43], spoilage microorganisms in beer [44], and for purifying distinct mammalian cells from a heterogenous pool [45–50]. Nonetheless, inertial microfluidic has not been applied to separation of neonatal cardiac cells.

While there are many available tools to measure cell size (like conventional imaging and counting or cell counter machines) and cell mechanotype (such as atomic force microscopy, micropipette aspiration, magnetic tweezers and optical stretchers) [51], these methods are not suitable for high-throughput measurement. To assess a large number of cells, we used a microfluidic-based method called Real-Time Deformability Cytometer (RT-DC). RT-DC is a contactless method that allows assessment of thousands of cells per minute [52–54]. RT-DC combines physical measurement and mechanical measurement in a single-step and permits the correlation of these properties in single cells.

The mathematical model used for RT-DC deformation calculation has been validated for measurement of non-spherical cells in multiple studies [55–58]. For example, RT-DC has been employed to characterise blood cells with different shapes: red blood cells (biconcave shape), white blood cells (irregular shape), control platelets (biconvex discoid) and activated platelets (octopus shape) [56]. RT-DC has also been employed to characterise cardiomyocytes that were derived from human induced pluripotent stem cells [59]. Indeed, RT-DC has been used to distinguish rod photoreceptor cells (rod-shaped) from other cells (e.g. cone shaped cells) in the heterogenous retina based on differences in morphology/size and mechanical properties [55].

Here, we describe a comprehensive method that permits the isolation, and physical and mechanical characterization of neonatal CMs in a rapid and high throughput manner. To this end, we exploited a simple sheathless inertial spiral microfluidic biochip to isolate primary mouse neonatal CMs from intact hearts. The biochip can be fabricated via cost-effective milling or 3D-printing methods. The mould fabricated by a micro-milling process does not require salinization. This method is cost effective and can be reused for extended periods of time. We also demonstrated that by using the same master mould, the device separation efficiency can be optimised by changing the location of the outlets (thus changing split flow ratio) by hole punching. This simple approach allows the user to carry out rapid optimisation or fine-tuning within their laboratory without the need to redesign and make a new master mould. Our isolation protocol did not negatively impact CM viability, growth or function. We also exploited RT-DC to phenotype the physico-mechanical properties of isolated CMs and showed that these cells have intrinsic biophysical properties that were distinct from non-CM cells in the heart. This study could pave the way for isolating minimally manipulated CMs for drug discovery and cell based therapy and for benchmarking properties of tissue engineered cells.

## 2. Results and Discussion

### 2.1. Label-Free CM Fractionation

It is known that CMs are significantly larger than their cellular counterparts in the heart [60, 61]. We investigated whether sorting cells based purely on their physical properties such as cell size, could help fractionate the heterogeneous cardiac cell population, and yield CMs that were intact, viable and able to proliferate.

To this end, we isolated neonatal hearts and generated single cell suspensions using enzymatic digestion (Figure 1A). The single cell populations obtained from digested hearts were suspended in 2% foetal calf serum (FCS) in phosphate buffered saline (PBS). These cells were then loaded into a syringe and introduced into the spiral microfluidic biochip using a syringe pump. The microfluidic biochip contains one inlet and two outlets (Figure 1B). The CMs were randomly dispersed in the suspension along with other cells. When these cells entered the microfluidic device, they gradually occupied the equilibrium positions. The equilibrium positions of cells and particles in the curved micro-channels are determined according to the Dean Flow Fractionation principle [62]. Based on this theory, particles flowing in a curved micro-channel, experience two major forces i.e. inertial lift forces (shear and wall induced lift forces), and Dean drag force [63, 64]. Lift forces are correlated with fluid density, fluid velocity, hydraulic diameter of the micro-channel and particle size. Drag force is correlated with dynamic viscosity of fluid and particle size. Interaction of these forces shifts the particle toward an equilibrium position, where the net force is zero. This position is mostly determined by particle size, when other parameters such as fluid density, viscosity, velocity and the micro-channel dimensions are constant [62, 65]. When applied to a heterogenous pool of cells, cells of different size focused at different positions in the cross-sectional plane of the micro-channel, which allowed size-based separation.

**Figure 1.**
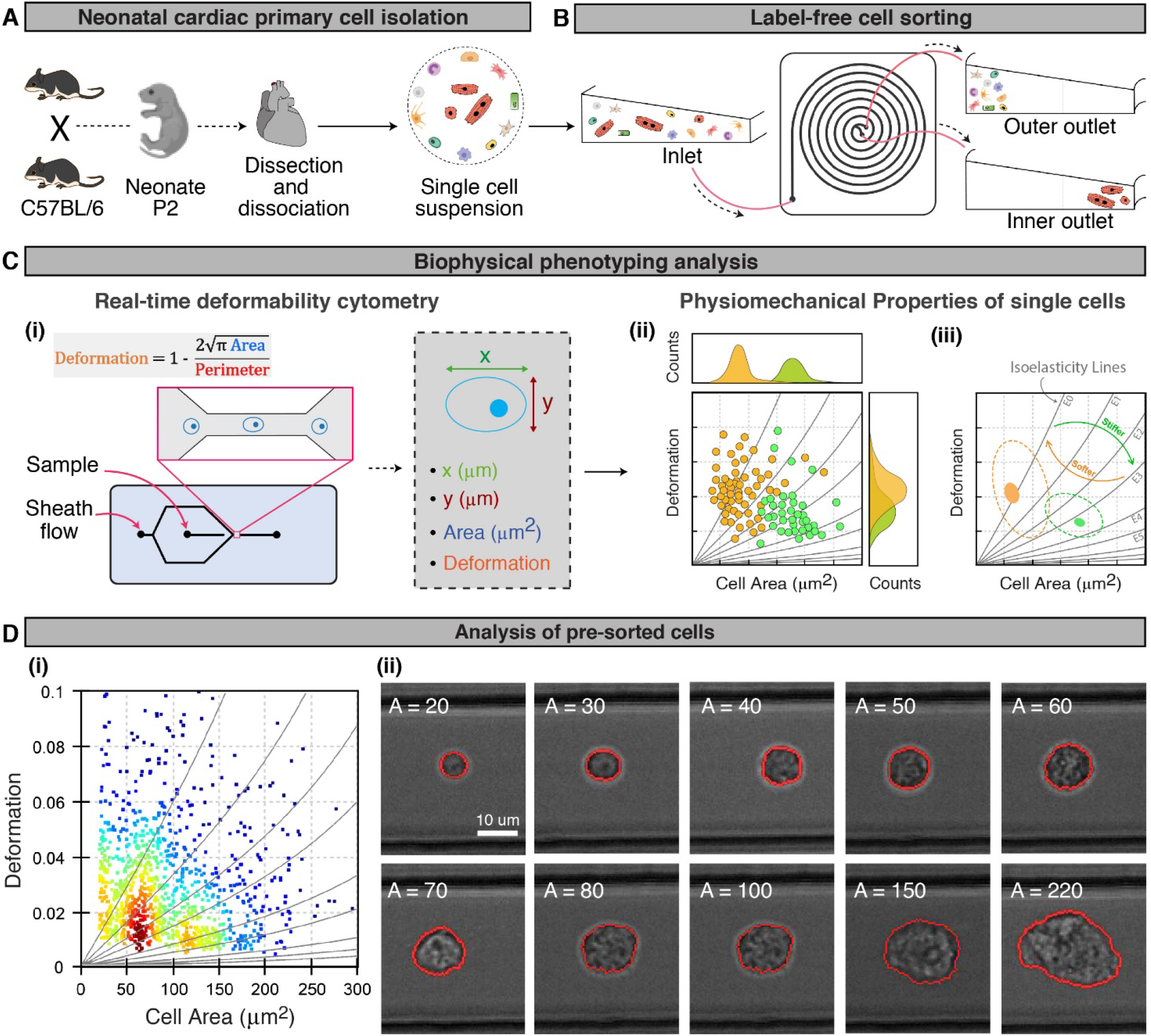
Overview of the experimental workflow. **(A)** Schematic outline of steps followed to isolate primary neonatal cardiac cell populations. **(B)** Primary cardiac cells are loaded into a spiral microfluidic biochip using a syringe pump, and cells were separated based on physical properties. **(C)** Schematic diagrams illustrate the flow of the analysis process. Physico-mechanical properties of sorted cells collected from the two outlets were analysed using real-time deformability cytometry (RT-DC). This method captures an image of every single cell that passes through the channel and measures physical (geometry, aspect ratio, cell size, etc.) and mechanical properties (deformation, Young modulus, etc.) of individual cells in real time. Results are represented as deformation vs. cell size scatter plots including isoelasticity lines, which divide a typical scatter plot into areas of identical stiffness. **(D)** Analysis of pre-sorted cells: scatter plot showing deformation vs. cell size of pre-sorted cells (left) and bright field images of single cells with a range of sizes (right) (A=20-300 μm^2^; any cell with size smaller than 20 μm^2^ and larger than 300 μm^2^ is considered as debris or bubbles. Scale bar=10 μm. (for more detail see Supplementary **Video S1**)

Next, to analyse the fractionated cells, we used a biophysical phenotyping approach-real time deformability cytometry (Figure 1C). Flow cytometry is a routine method that uses cell diameter as an index for comparing cell size. However, CMs have a rectangular shape in adult hearts and are not completely ovoid at the embryonic or neonatal stage. Previous studies have used a parameter termed the equivalent spherical diameter (ESD) to describe irregular shape objects [43]. Whilst using conventional microscopic imaging combined with image analysis is a time-consuming process, conventional cytometers are imprecise given that they only report a few basic parameters such as diameter and cell number. Conventional cytometers also rely on matching measurements to pre-defined cell types (e.g. fibroblasts or blood cells). Furthermore, none of these methods provide rapid, high-throughput and comprehensive analyses.

We used a microfluidic-based system that was attached to an epifluorescence microscope and a high-speed camera that enabled real time study of sorted cells directly after processing [52–54]. The workflow of the analytical steps is illustrated in Figure 1C. In this system, a queue of cells passes through a straight microchannel while a high-speed camera takes an image of every single cell, and measures the dimension of those individual cell (herein referred to as x, and y). Based on these measurements, an algorithm calculates geometric indices such as perimeter, cell area, circularity and aspect ratio (Figure 1C(i)). Moreover, while cells were experiencing hydrodynamic shear and normal stresses, the extent of deformation of cells was evaluated using an image-processing algorithm [53]. Therefore, this method can be utilised to analyse the mechanical properties of cells (e.g. stiffness or young modulus (E)). These data can be cumulatively represented as a scatter plot (Figure 1C(ii)) or distribution histogram (Figure S1), and distinct cell types can be categorised by contour plots (Figure 1C(iii)) [51].

We analysed neonatal cardiac cells before and after sorting (Figure 1D (i) and Figure S1). Distribution of cell sizes based on x and y are different, so we considered cell area that includes both dimensions as a suitable criterion of cell size. Hereafter, all results reported in this study were based on cell area (μm^2^). The scatter plot depicts cell deformability vs. cell size, while the curved grey lines are the isoelasticity lines (Figure 1C (ii) and (iii)), as a measure for elasticity and stiffness of the cell [54]. Moreover, the bright-field image of every cell can be captured for further analysis (Figure 1D(ii)). We performed CM isolation using a widely used method, i.e. magnetic activated cell sorter (MACS) and measured CM size using a haemocytometer (data not shown here).

Neonatal CMs have an average area (A) of higher than 100 μm^2^ as measured in our preliminary experiments (Figure S2, supporting information). Thus, we attempted to fractionate the cardiac cell population into two separate groups: A≥100 μm^2^ and A<100 μm^2^ as CMs and non-CMs respectively. It was also noted that the nuclear morphology of cells could be visualised from the bright-field images, in which each cell can be identified as either a mono-nucleated cell (with a single small round nucleus) or as multi-nucleated cell (with large irregular shaped nuclei) (Figure 1D(ii)).

### 2.2. Flow Rate Optimisation Based on Biophysical Properties

The spiral microfluidic biochip we used in this work has a trapezoidal cross section (600 μm channel width, with inner and outer wall height of 80 and 130 μm respectively). The asymmetry of the trapezoid cross-section leads to higher focusing of particles and better separation resolution compared with prior designs [39, 62]. As flow rate (Q) is a determinant of separation performance, this microfluidic design has been deployed to isolate particles and cells of various sizes e.g. polystyrene microbeads [62], circulating tumour cells (CTCs) [46], mesenchymal stem cells (MSCs) [48], blood plasma [40], chondrocytes [50], and to separate debris and dead cells from live cells in bioreactors [66]. The optimal processing flow rates for specific size ranges have therefore been calculated. Based on these studies and our preliminary experiments using mouse MSCs and adult CMs, we tested four flow rates (e.g. 1.0, 1.2, 1.4, and 1.6 ml/min) to determine the optimal flow condition for CM separation. Flow rates below and above these magnitudes were not tested as cells will mainly migrate to one side of the microchannel. Processed samples collected from the outlets were subject to Real-Time Deformability Cytometry (RT-DC) analysis (Figure 2A). Results comparing different flow rate conditions showed that large cells were dominant in the inner outlet when Q=1.2 ml/min. However, some large cells were not fully captured through the inner outlet and were mixed with smaller cells in the outer outlet (Figure 2B). The cell area profiles of collected cells in Figure 2C showed a big peak of large cells (A≥100) at Q=1.2 ml/min. In addition, the cell area profiles between inner and outer outlet were most distinct at Q = 1.2 ml/min when compared to other flowrates being tested. Increasing the flow rate to 1.4 and 1.6 ml/min resulted in more small cells in the inner outlet which was contrary to our aim. Hence, we considered Q=1.2 ml/min as the optimal flow rate to capture CMs using a serial sorting strategy, where cells going to the outer outlet could be further separated by a repeat round of sorting. As illustrated in the histograms in Figure 2D, the deformation profiles of cells from the inner and outer outlets were similar for all flow rates tested except for Q = 1.2 ml/min. There is a shift in cell area peaks and deformation peaks at Q = 1.2 ml/min, indicating that there are differences in both cell size and cell stiffness between the inner and outer outlets.

**Figure 2.**
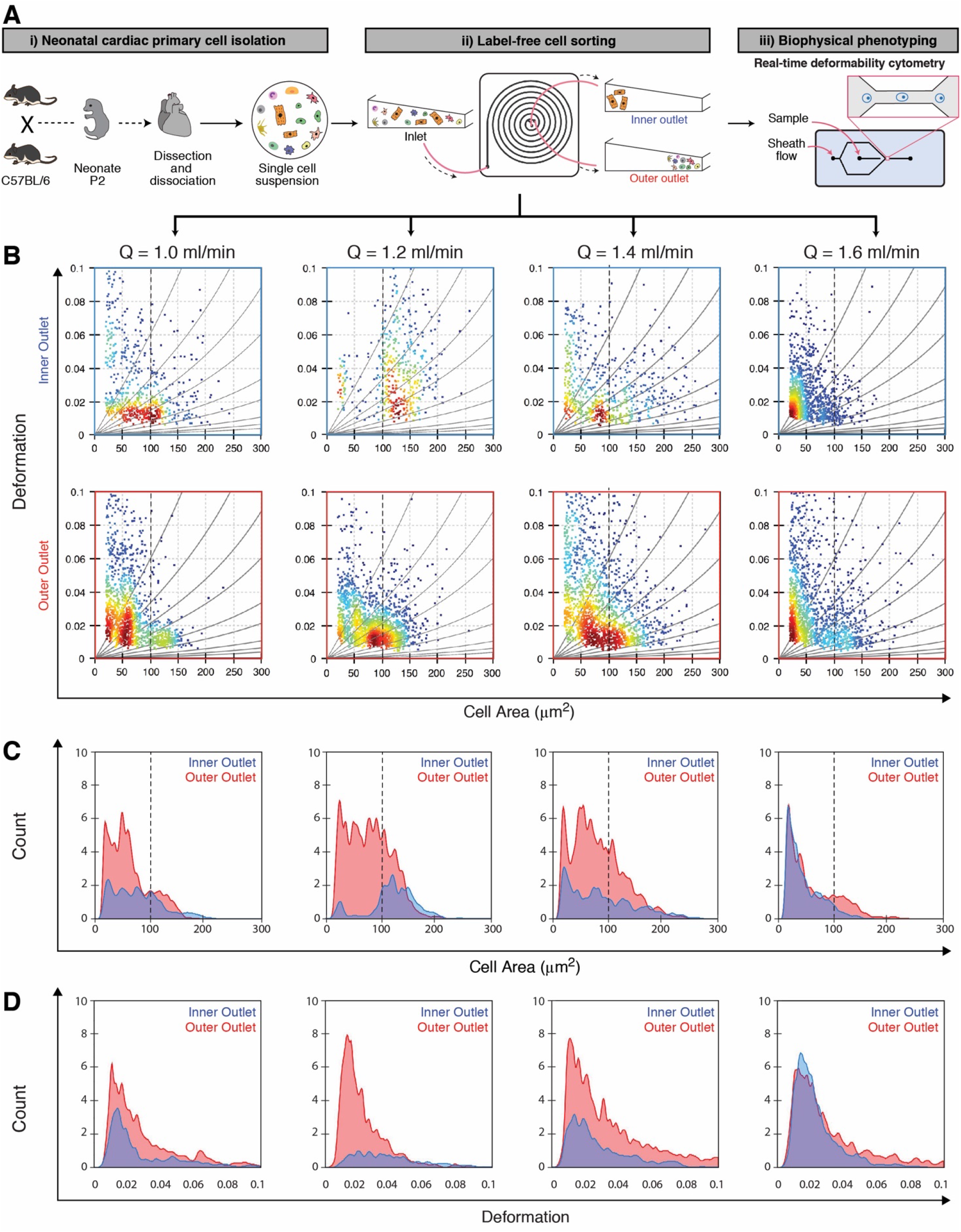
Label-free sorting of neonatal cardiac cells followed by biophysical phenotyping analysis shows optimal separation at Q=1.2 ml/min. **(A)** Schematic representation of experimental design: i) Neonatal cardiac cell isolation, ii) cardiac cells loaded into microfluidic setup via different flow rates (Q = 1.0, 1.2, 1.4, 1.6 ml/min) and collected from the two outlets and iii) collected cells were analysed using the RT-DC method. **(B)** Scatter plots of cell size vs. deformation of the sorted cells. **(C)** Overlapped histograms of cell size distribution of sorted cells collected from the spiral device. Blue histograms and red histograms showing area of cells collected from the inner outlet and outer outlet, respectively, at different flow rates. At Q=1.2 ml/min there was a significant peak showing large cells (A>100) were separated from smaller cells collected from the inner outlet but some residual large cells remain in the outer outlet. **(D)** Overlapped histograms showing the deformation values of processed cells collected from the inner outlet (blue) and outer outlet (red) at different flow rates.

To verify if the deformation profiles reflected the intrinsic mechanical properties of sorted cells or if the change in deformation profiles were attributed to damage caused by the sorting process, mechanical phenotyping was performed on pre-sorted cells (untreated control). We observed that cells in the inner and outer outlet together displayed a similar range of deformation values as pre-sorted cells, regardless of the flow rates (Figure S4 in supporting information). These results indicated that cells passing through the spiral microchannel were not damaged by shear stress. These results also confirmed that the cells collected from the inner outlet were both larger and stiffer compared to those collected from the outer outlet at Q = 1.2 ml/min. In addition, all CMs and non-CMs collected from spiral microfluidic sorting showed high viability (>95%, determined by trypan blue staining), comparable to previous reports for separation of CTCs, MSCs, and chondrocytes [45, 46, 50].

Analysis of mechanical properties of cardiac cells is important and a few recent studies have shown that changes in mechanical properties of cells is a biomarker of cell state and can be used as a label-free diagnostic tool for diseased cells [67, 68]. For instance, it has been shown that changes in the mechanical fingerprint of specific cell populations is linked to disease progression such as malignant pleural effusions [69], graft-versus-host disease (GVHD) [70], and blood-related diseases such as leukaemia, malaria, and bacterial and viral infections [71].

### 2.3. Tunability of the Split Flow Ratio to Optimise Separation Performance

Apart from flow rate, there are other parameters that can impact the separation performance of a microfluidic device [66]. One is flow split ratio (SP) which is defined as the ratio of fluid coming out from the outer and inlet outlet per time unit. Flow SP can be modulated by changing the tubing length of outlets, diameter of tubes connected to the outlet, and the location of tubes in the outlets. Therefore, we investigated the possibility of improving the separation performance by optimising the device SP. Three devices were microfabricated as shown in Figure 3A-D using the same master aluminium mould that was designed with extra-long branching tubes. Outlets were punched on locations along the branching tubes of each device that were different from the other two devices (Figure 3C), such that the channel length of the outlets were different in each of the three devices as shown in Figure 3E-F. In this manner, the SP can be easily tuned without the need to redesign and to mould a new master. The flow split ratio of the three devices SP1, SP2 and SP3 was measured and determined to be 2.4, 2.0 and 1.1, respectively (Table S1, supporting information).

**Figure 3.**
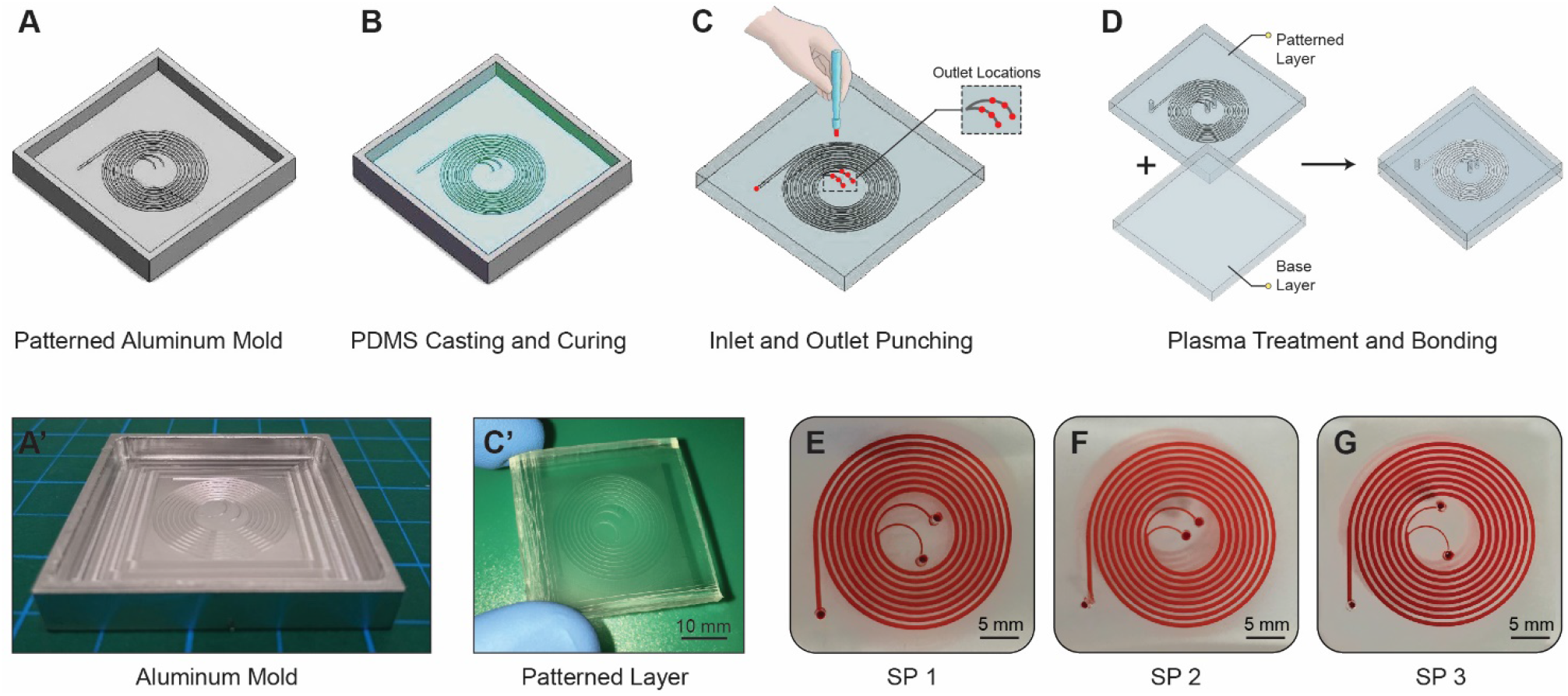
Schematics showing the workflow of microfabricating three devices each with a different flow split ratio (SP) using the same master mould. **(A)** The master aluminium mould was designed with extra-long branching tubes. **(A’)** Photograph of an aluminum master mould having the microfeatures of spiral biochip, the mould was microfabricated by micromilling. **(B)** PDMS was cast onto the aluminum mould and cured using standard soft-lithography technique. **(C)** The PDMS layer containing the microchannel pattern was detached from the master. Outlets were punched on selected locations along the branching tubes in order to control the channel length of the outlets. **(C’)** Photograph showing a PDMS substrate with spiral microchannel pattern. **(D)** The micropatterned PDMS layer was plasma treated and bonded to a base layer PDMS slab to form the spiral device. **(E-G)** Photos of three microfluidic devices each with different outlet locations. The difference in outlet locations gives rise to different outlet tube length, thus different SP. The microfluidic devices were filled with red food dye for visualization purpose.

We applied the same experimental design, the same RT-DC measurement, and used the optimal flow rate (Q = 1.2 ml/min) to test the performance of the three devices with different split flow ratio (Figure 4A). First, the effect of split flow ratio on size distribution of cells collected from the outlets was evaluated for the three devices. The overlapped histograms in Figure 4B showed that device SP2 displayed more distinct peaks when comparing its cell area profile in the inner outlet to that of its outer outlets. Whereas using the other two split ratios led to less large cell enrichment in the inner outlet and higher numbers of large cells mixed with small cells in the outer outlet. We further compared the separation performance of the three devices by comparing their large cell recovery percentage. The recovery percentage was calculated as the number of large cells collected in an outlet over the sum of large cells present in both inner and outer outlet of that given device. As shown in Figure 4C, device SP2 yield the highest large cell recovery rate, in which approximately 70% of large cells were recovered in the inner outlet, making it the most efficient device amongst the three.

**Figure 4.**
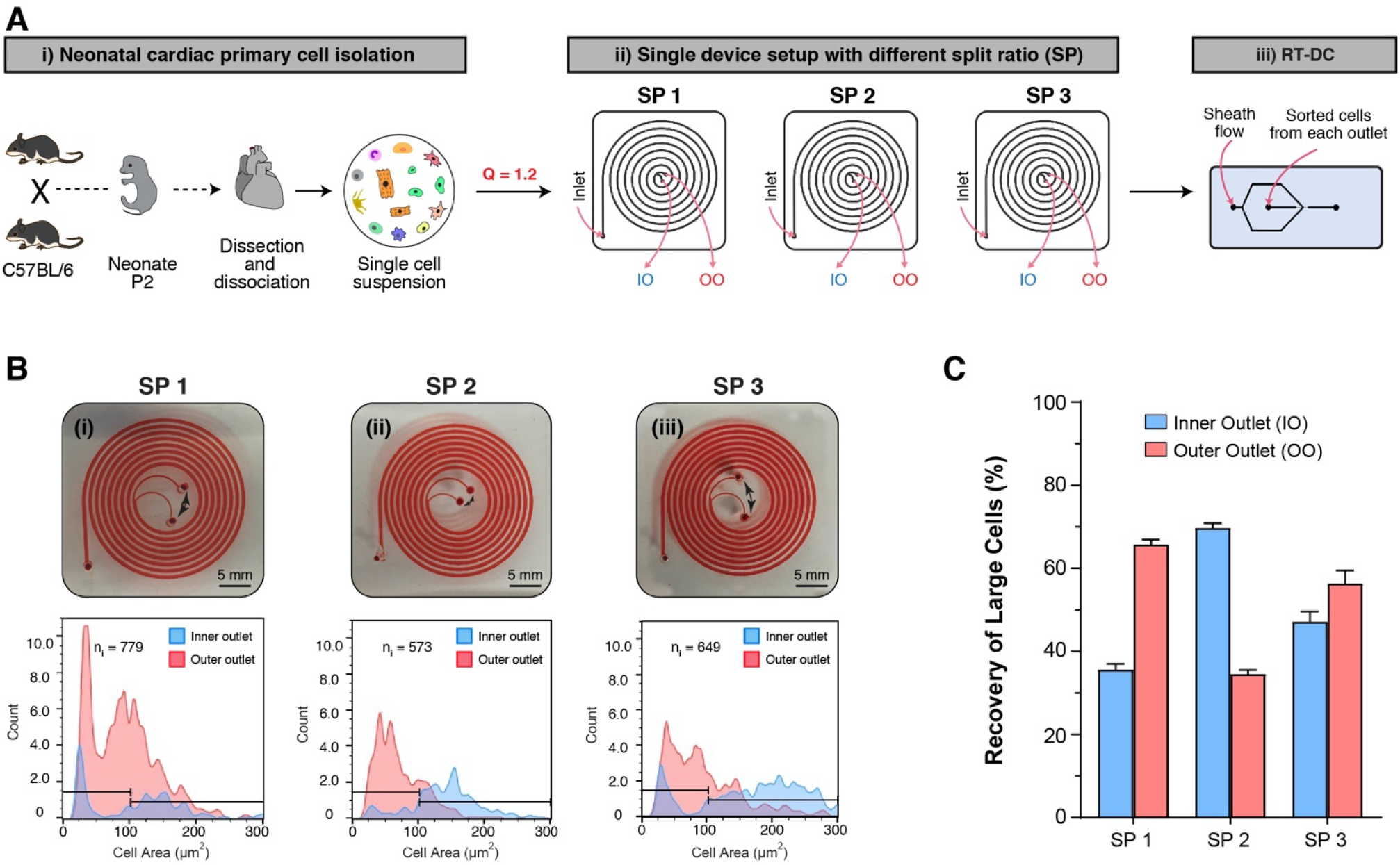
Improving the separation performance by changing the split ratio (SP). **(A)** Schematic representation of the experimental design: i) Neonatal cardiac cell were isolated, ii) cardiac cells loaded into 3 microfluidic devices each with a different flow split ratio (SP1 = 2.4 SP2 = 2.0, SP3 = 1.1). The cells were fed at an optimal flow rate of Q=1.2 ml/min. iii) Cells collected from each outlet were analysed using the RT-DC method. **(B)** Photos of three microfluidic devices with different SP. The SP was modulated by changing the outlet tube length thus changing fluidic resistance. The microfluidic devices were filled with red food dye for visualization purpose. Overlapped histograms showing the effect of change in split flow ratio on size distribution of cells in inner and outer outlets. Blue histograms and red histograms showing the size of cells (in cell area) collected from the inner outlet and outer outlet, respectively. The left gates show the cells with cell area smaller than 100 μm^2^, while the right gates show the cells with cell area larger than 100 μm^2^. **(C)** Bar chart comparing the percentage of recovery of large cells in the inner outlet and outer outlet for a given SP.

### 2.4. Cascaded Configuration for Improved CM Recovery

A cascaded setup with two consecutive devices was utilized to improve large cell recovery efficiency. Two consecutive devices were connected through the outer outlet of device 1 and the inlet of device 2 (Figure 5A). Cardiac cells were first sorted in device 1 and cells from the first outer outlet were passed to the inlet of device 2. Both devices in the cascade setup were operated using optimised conditions (flow rate of 1.2 ml/min and SP 2). Large cells were collected from the inner outlets of both devices 1 and 2, while smaller cells were collected in the outer outlet of device 2. Sorted cells were subjected to RT-DC mechano-phenotyping. Using RT-DC, we captured high-speed microscopic video images. As shown in the representative images in Figure 5B and Figure S3 (supporting information), the majority of cells collected from the inner outlets were cells with larger cell area, while the majority of cells collected from the outer outlet were smaller mononuclear cells. However, there was a small population of small non-CM cells (9.8% – determined by FlowJo) in the inner outlet (Figure 5B(i)). This was expected, given the overlap in size between immature CMs (i.e. mononucleated) with other mononuclear cardiac stromal cells [72]. It is common for occasional small cells to appear with large cells in the inner outlet channel as seen in other spiral inertial studies [39, 62, 73, 74]. This phenomenon has been investigated and described by Guzniczak et al, [75]. In an inertial focusing channel, cells/particles have the tendency to form trains with defined inter-particle spacing. The inter-particle spacing is believed to result from particle-induced convection, and the inter-particle spacing is sustained by inertial lift force. It is known that particle interactions especially particles in a heterogeneous mixture, can shape and alter the equilibrium position and particle spacing, thus altering the fluid flow pattern in inertial systems. It has been reported that small particles can occasionally deviate from the equilibrium position and self-assemble into the train of larger particles, resulting in a change in separation efficiency [75]. The hydrodynamic mechanism involved in deviation in particle focusing positions remained poorly understood [74, 75].

**Figure 5.**
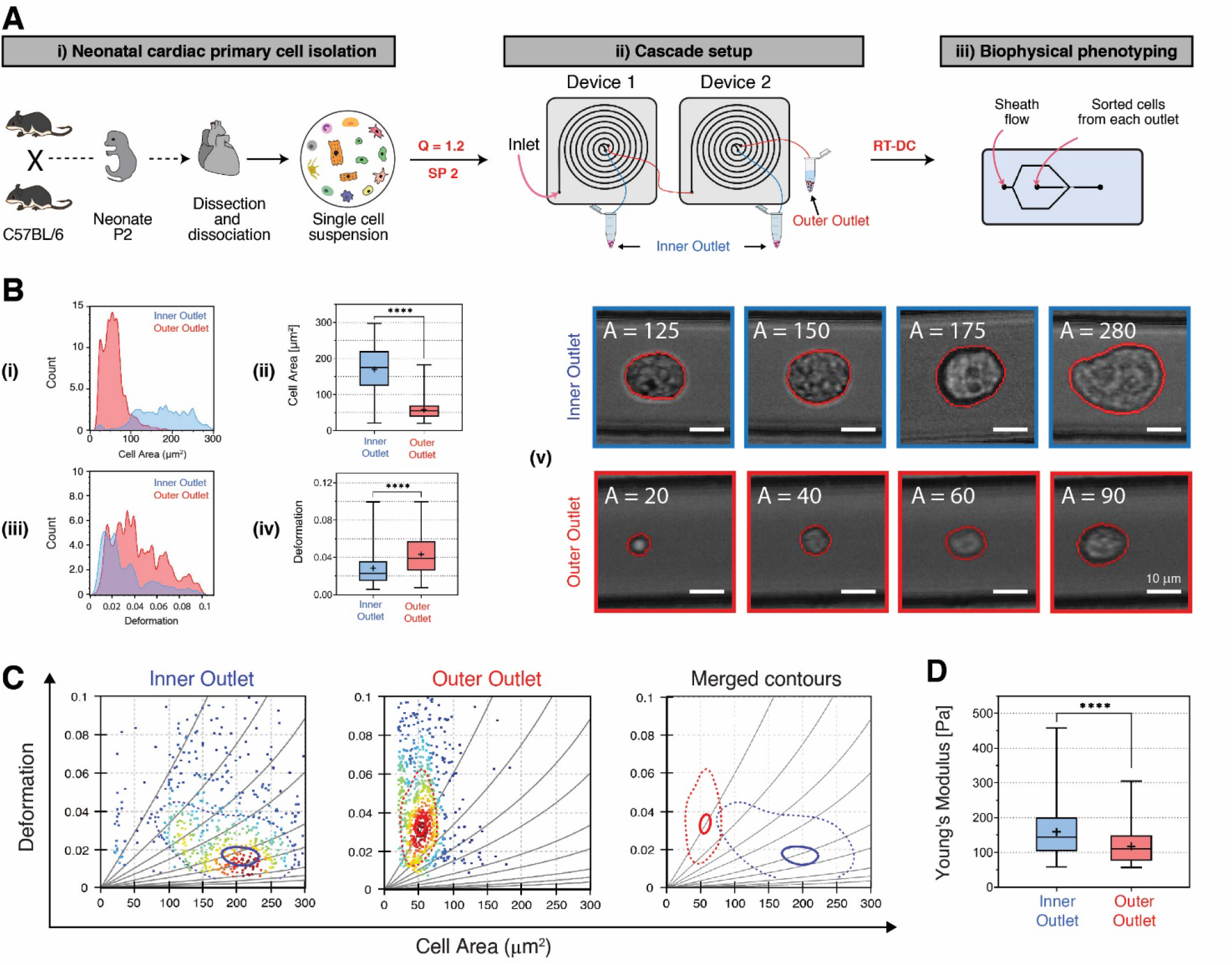
Isolation of large cells from the whole cardiac cell population in a 2-stage cascaded microfluidic system. **(A)** Schematic representation of the protocol including i) cardiac cell isolation, ii) 2 stage label-free cell sorting operated at optimal condition (Q=1.2 ml/min and SP2). Large cells were collected from the inner outlets of both device 1 and 2, while small cells were collected from device 2 only. iii) Biophysical phenotyping analysis using RT-DC. **(B)** (i) Overlapped histograms representing size distribution of sorted cells in the inner outlets (blue) and outer outlets (red). (ii) Box and whisker plot show a significant difference in cell size between populations collected from the inner outlets and outer outlets (iii) Overlapped histograms representing deformation distribution and of sorted cells. (iv) Box plot shows a significant difference in deformation value between cells collected from the inner outlets and outer outlets. (v) Representative images highlighting large cells collected from the inner outlets (blue box) and small cells collected from the outer outlets (red box) of the cascade system; for more detail see Supplementary **Videos S3 and S4**. Scale bar: 10 μm. **(C)** Deformation versus cell area scatter plots with 95%-density (dashed lines) and 50%-density (solid lines) contour plots. Blue contour gates and red contour gates highlight the cell events from the inner outlets and outer outlets, respectively. Most of the cells from the inner outlets (gated by blue contours) are stiffer than most of cells from the outer outlet (gated by red contours). **(D)** Box and whisker plot shows a significant difference in Young Modulus for the sorted cells populations between the outlets (**** *p*<0.0001). For all box and whisker plots, solid line within the box denote the median extending from the 25th to 75th percentiles and error bars span minimum to maximum values within the indicated datasets.

It has previously been noted that this setup is amenable to high throughput analysis (50,000 cells/ml × 1.2 ml/min × 2 chip = 120,000 cells/min) with throughputs up to 1 L/min achieved by parallelization of biochips via multiplexed or stacked designs [35, 38, 40, 46]. The RT-DC algorithm used an analytical model and numerical simulation [52, 53] to provide isoelastic lines that relates cells of different sizes to their stiffness values. Hence mechanical fingerprints of cell populations can be explored using deformation-cell size plots. The contour plots analysis (Figure 5C) showed that smaller cells have lower E values (pliant phenotype), whereas the contour of the enriched large cell population expressed higher E values (stiffer phenotype). The enriched population of CMs contained mainly larger cells which were stiffer in nature (the population that are less deformed in constriction; blue contours; Figure 5C-left and right), while mono nucleated cells are smaller and pliant in nature – red contours (Figure 5C-middle and right). Figure 5D revealed that the Young Modulus of cells collected from the inner outlet were significantly higher than the Young Modulus of cells collected from the outer outlet. This result further confirmed that the cells from the inner outlet were stiffer compared to the cells from the outer outlet.

The optimisation steps taken to improve separation performance can be summarised as follows: in early experiments, a single spiral device design (SP1, split flow ratio = 2.4) was employed and the separation performance was optimised by adjusting the flow rate to Q2 = 1.2ml/min. Although more large cells were collected in the inner outlet at Q2 as compared to other flow rates being tested, there was a size overlap between yields from the inner and outer outlet at Q2 as shown in the size distribution profile in Figure 6A. Subsequently, separation efficiency was improved by modulating the split flow ratio (SP2 design) with the flow rate fixed at Q2 (Figure 6B). Separation efficiency was further improved by using a 2-stage cascade setup in which two spiral devices were connected and operated in parallel. As shown in Figure 6C, a larger difference in size distribution between the inner and outer outlet was obtained using a cascaded setup (Q2 & SP2). Figure 6D showed the incremental improvement in separation efficiency, in terms of large cell recovery rate, using a combination of optimisation methods. It is also worth mentioning that our approach allows CMs to be isolated in a rapid and high throughput manner (2.4×10^5^ cells/min). The flow rate used here (1.2 ml/min) was 15 to 60 times faster than other reported inertial methods [10, 11].

**Figure 6.**
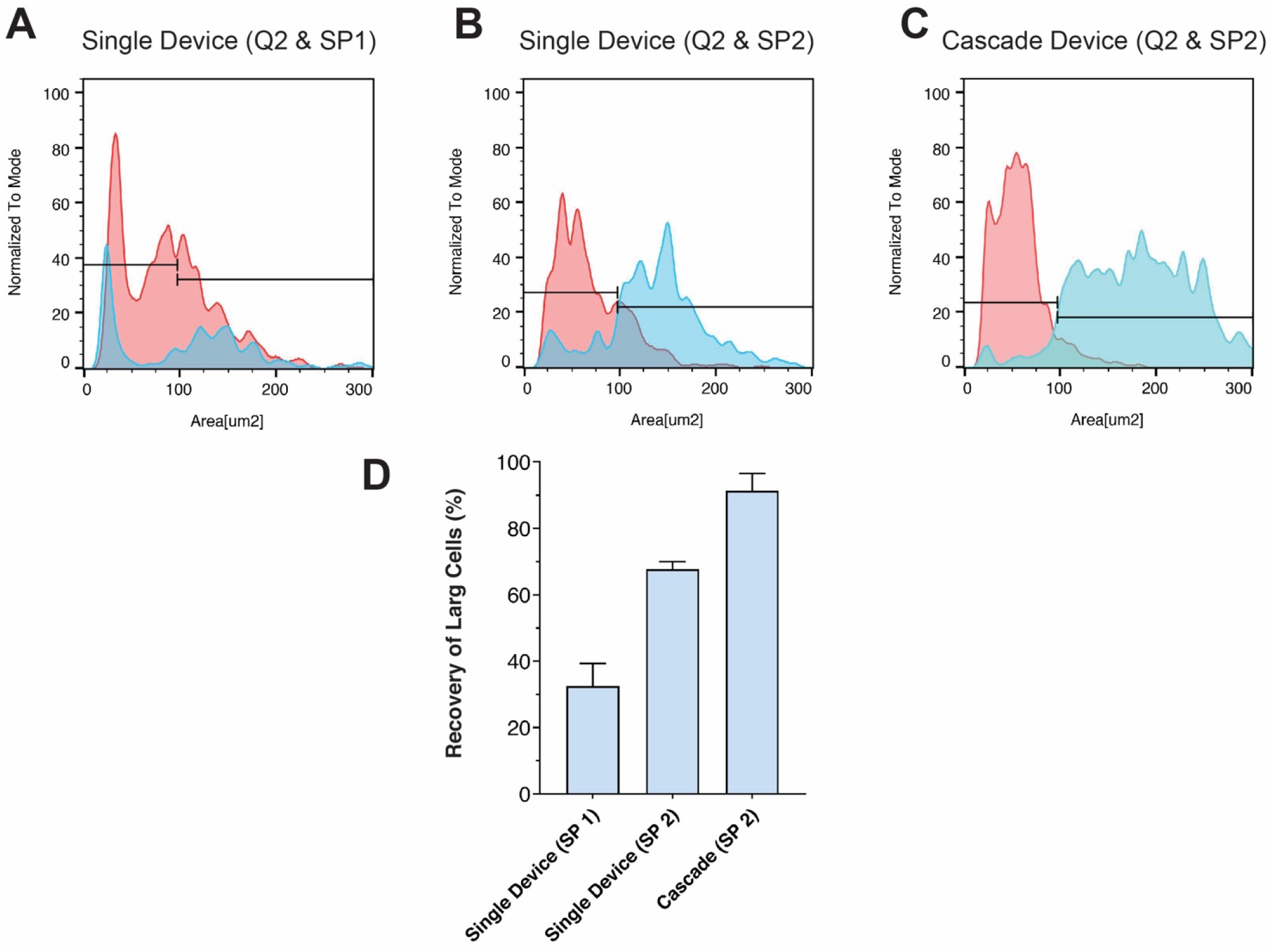
Incremental improvement in separation performance. **(A)** The spiral microfluidic device was first optimised by adjusting its inlet flow rate. An inlet flow rate Q2 = 1.2 ml/min was found to be the optimal flow rate. **(B)** The device was further optimised by modulating its split flow ratio to 2 (SP2) while fixing the inlet flow rate at Q2. **(C)** Finally, the separation performance was further improved by using a cascaded configuration with two connecting devices operating at Q2 and SP2. Overlapped histograms show the size distribution of cells in inner outlets (blue) and outer outlet (red) from three different configurations. **(D)** Bar chart showing an incremental improvement in separation efficiency, in terms of large cell recovery rate, by the use of a combination of optimisation methods.

### 2.5. Characterization of Recovered CMs by Immunostaining

To investigate whether this method works in different mouse strains, we used two distinct strains of mice for CM extraction- (i) C57BL/6 (wild type (WT)) and (ii) *Gt(ROSA)26Sor^tm14(CAG-tdTomato)Hze^* (tdTomato) transgenic mouse strains. In each experiment, microfluidic sorting was performed using a spiral biochip with a cascaded configuration operated at Q = 1.2 ml/min and SP2 (Figure 7A(i) and B(i)), the sorted cells were analysed using RT-DC (Figure 7A(ii) and B(ii)). At the end of cell sorting, the sorted cells were plated and cultured on matrigel coated dishes for 48 hrs (Figure 7A(iii) and 5B(iii)). At the end of 48 hrs, we assessed the phenotypic and functionality of these cells using confocal microscopy and live cell imaging.

**Figure 7.**
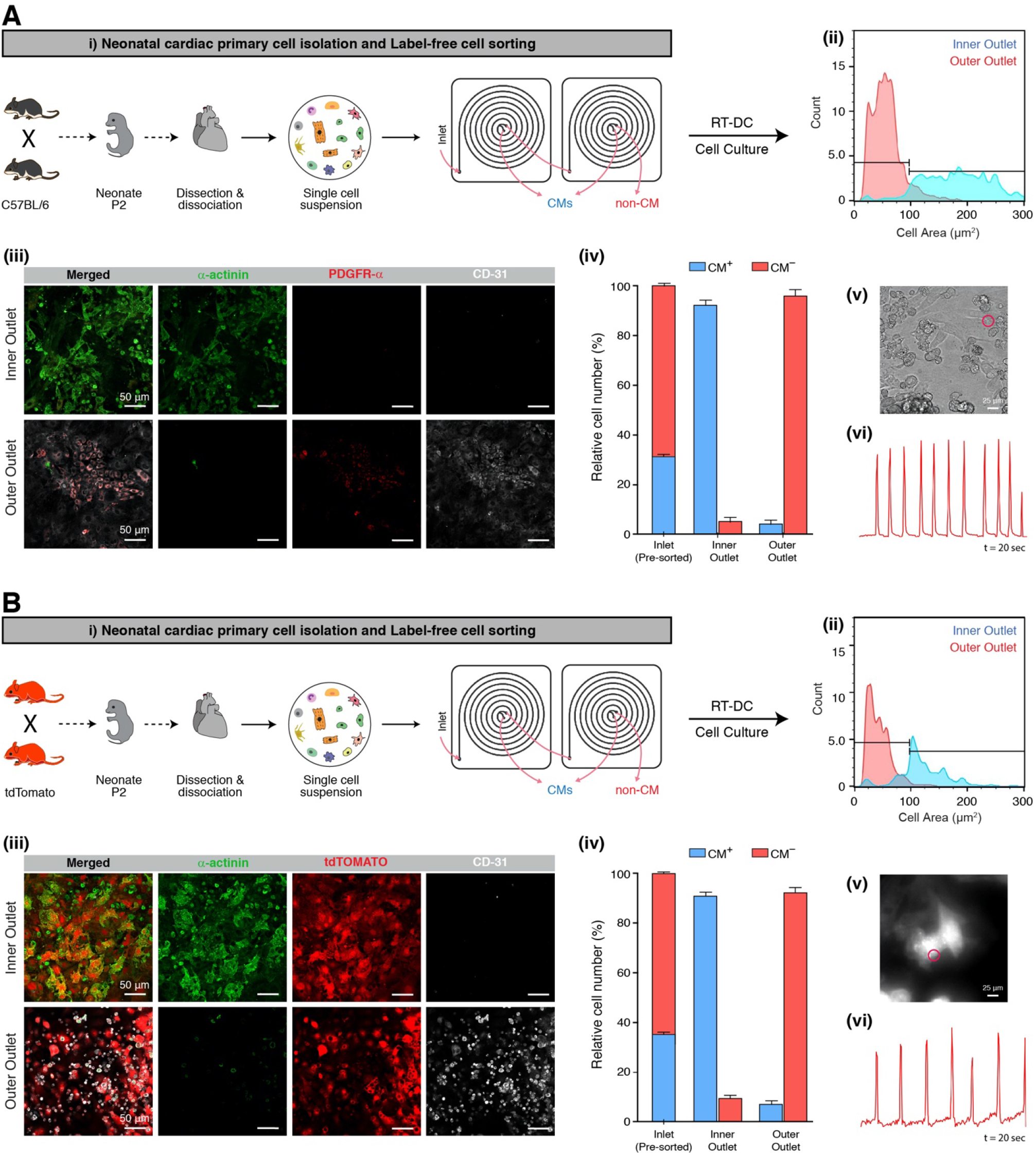
Characterization of neonatal CMs sorting for two different mice type A) C57BL/6 WT and B) tdTomato transgenic mice. **(i)** Schematic representation of the protocol of cardiac cell isolation followed by label-free cell sorting. **(ii)** Histogram showing large cells separation in inner outlet (blue profile) from small cells in outer outlet (red profile). **(iii)** Confocal fluorescent microscopy images of sorted cells into inner and outer outlets for CM cell marker α-actinin (green), CD-31 (endothelial; grey), and other cardiac cells markers PDGFRα in (A) C57BL/6 wild-type mouse or (B) tdTomato (stromal; red) in (B) *Gt(ROSA)26Sor^tm14(CAG-tdTomato)Hze^* transgenic mouse. Scale bars: 50 μm. **(iv)** The purity of cells collected from each outlet were quantified by counting the percentage (or relative cell number) of α-actinin expressed CM (CM+) and α-actinin negative mono-nuclear cells (CM-) collected from pre-sorted (before sorting control), inner outlet and outer outlet. **(v)** Still images of beating CMs after 2 days of culture and **(vi)** plot showing the beating signals over time. Scale bars: 25 μm.

The cells were immunolabelled prior to examination using confocal microscopy. Representative confocal microscopy images in Figure 7A(iii) and 7B(iii) showed that the cardiac specific marker (cardiac α-actinin) was highly expressed in cells were collected from the inner outlet. Cells collected from outer outlet showed expression of endothelial (CD31) and stromal (PDGFRα) markers but expressed very few cardiac α-actinin markers. These results indicated that there was a significant enrichment of CMs amongst cells collected from the inner outlet for both WT and tdTomato strains, while cells in the outer outlet were enriched for vascular and stromal mono-nuclear cells.

The separation efficiency was evaluated by quantifying the purity of cells collected from inner and outer outlets for both mouse strains. This was performed by counting the number of immunolabeled cells in both outlets to calculate the percentage of CMs and non-CMs (or relative cell number) in each group (Figure 7A(iv) and B(iv)). For wild type mouse, 90.66 ± 2.16 % of the cells in inner outlet were α-actinin expressing cardiomyocyte (CM+), while 95.67 ± 2.92 % of the cells in the outer outlet were of CD31 or PDGFRα expressing non-CM cells (CM-). For tdTomato transgenic mouse, 90.45 ± 1.23 % of the cells in inner outlet were α-actinin expressing cardiomyocytes (CM+), while 91.09 ± 2.16 % of the cells in the outer outlet were α-actinin negative non-CM cells (CM-). These results further confirmed that the larger and stiffer cells collected from inner outlets of the devices were primarily CMs.

The small difference in separation efficiency between wild type and transgenic mice, as reflected by the small difference in percentage of CM and non-CM in Figure 5A and 5B, was most likely due to the difference in cell size between the two strains. Preliminary data (Figure S7 supporting information) showed that there was some difference in CM cell size between C57BL/6 wild type and tdTomato transgenic mice. This observation was not surprising, as the cell sizes of CMs are known to increase during the postnatal growth period. Leu et al., have reported that CMs from knockout mice were characterised by a high variability in cell size as compared to wild type mice [76]. Indeed, cell morphology including cell size, varies from mouse-to-mouse within the same strain [77]. This further highlighted the functionality of our sorting approach that allows users to carry out rapid optimisation without the need to redesign a new master mould, thus enabling users to easily adjust sorting parameters based on the cardiac cell size range of their animal species of choice.

CMs account for approximately 30% and 35% of the total number of cardiac cells in wild type and transgenic neonatal mouse hearts, respectively (Figure 7 A(iv) and B(iv)). Highly purified CMs are desirable for use in disease model systems, drug testing and cell therapy applications in regenerative medicine, as impure or mixed cell populations may interfere with the function of CMs [78]. Therefore, it is important to optimize efforts to remove small cells in order to obtain CMs with high purity. Herein, we demonstrated that by optimising flow rate, split flow ratio and using a cascaded configuration, we were able to isolate neonatal CMs with >90% purity.

As expected, the purity of sorted cells was not 100%, which could be due to size overlap between different cell types. Cells that were cultured likely underwent proliferation, which also contributes to discrepancies in cell size (as cells displayed different size at different growth phase). In addition, the deviation in particle focusing position as discussed earlier, may also lead to a lower purity in the inner outlet. The separation performance of our system is comparable to other published inertial microfluidic systems [79]. For example, Mach & Carlo used a straight channel inertial microfluidic system and were able to remove 80% of pathogenic bacteria from volunteer blood samples [80]. Che et al., used a contraction expansion array channel (CEA channel) inertial microfluidic system to enrich rare circulating tumor cells from clinical blood samples with a capture efficiency of 83% [81]. On the other hand, Lee et al., used a CEA inertial microfluidics device to separate cancer cells from samples prepared by spiking breast cancer cells into diluted whole blood, obtaining a recovery rate as high as 99.1% [82]. Lei & Dandy have reported the use of a sinusoidal channel inertial microfluidics system to separate cyanobacteria from samples that contained pure lab-grown cyanobacterium PCC6803 in culture medium, and were able to enrich PCC6803 with a recovery efficiency of 98.4% [83]. A detailed review of the performance of various microfluidic devices can be found in Gou et al. [79]. It is worth mentioning that when comparing separation performance, one should also take into account the composition factor. The focusing behaviour of an inertial microfluidic system is highly dependent on the composition of the feed with mixed populations known to yield lower separation efficiency compared to pure populations [75].

Finally, we assessed the beating behaviour of enriched CMs using a spinning disk microscope (Figure 7A(v) and B(v)). The CMs isolated from both WT and tdTomato mice maintained their contractile ability and showed similar contraction patterns over time (Figure 7A(vi) and B(vi) and Supplementary videos S4 and S5).

In summary, CMs are the focus of cardiac regenerative studies. They switch from mono-nucleate to mature bi/multi-nucleate cells which exit the cell cycle and stop proliferating. Many studies have focused on CM maturation [84–86]. However, there is no agreement regarding the structural, functional and metabolic factors that drive this process. Factors such as cell cycle activity, size, age, polyploidization/multi-nucleation, number of nuclei per CM impact post-injury cardiac function. Neonatal CMs, which have a high regenerative capacity are an invaluable source for addressing some of these questions. We have developed a simple spiral microfluidic biochip-based method to isolate mouse neonatal CMs. This method isolates and efficiently separates mono-, bi- and multinucleated CMs from mononuclear stromal cells in the neonatal mouse heart. This method of CM isolation depends on flow rate, which is determined by the biophysical properties of cells. Confocal fluorescent microscopy analysis of isolated CMs (Figure 8) indicated that enriched cells retain key features of CMs; (i) immature or mono-nucleated cells that mainly exist in the embryonic heart. These cells were smaller than other CMs and have indistinct intercalated disc striation in microscopic images (Figure 8(i)), (ii) Partially mature but still mono-nucleated cells that possess the highest proliferative capacity (Figure 8(ii)), and (iii) mature CMs that are bi-nucleated, large and non-proliferative cells (Figure 8(iii)). CMs that were isolated using this device showed contractile properties in keeping with their identity.

**Figure 8.**
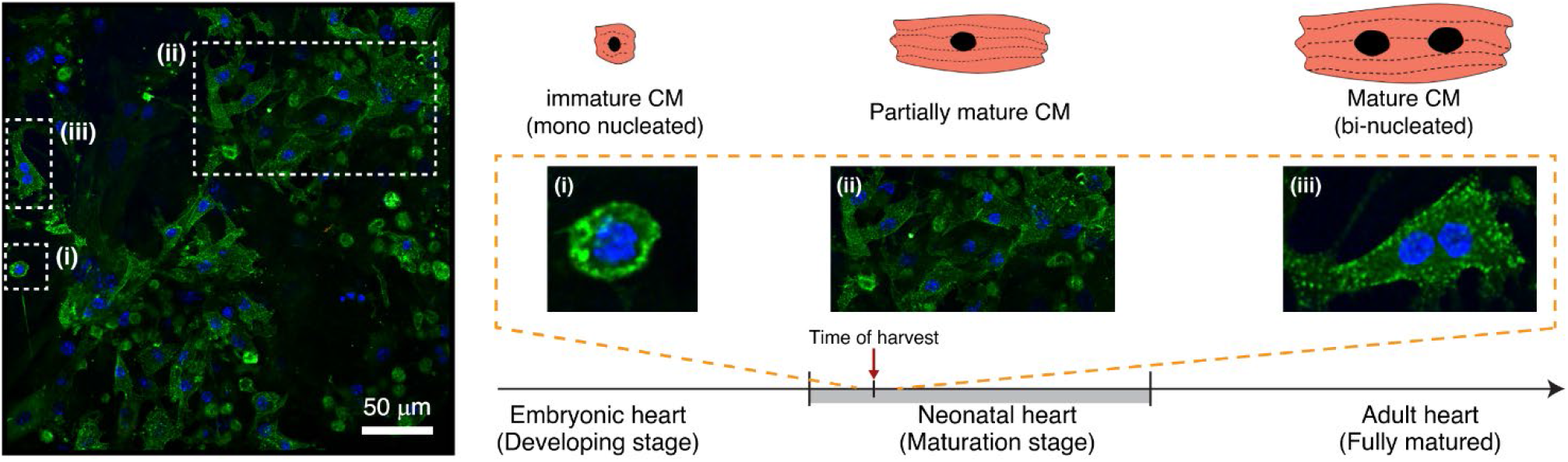
Confocal fluorescent microscopy images of enriched CMs isolated from P2 mice having three types of CMs. **(i)** immature and mononucleated CMs. **(ii)** Partially mature and mononucleated CMs. **(iii)** Mature and binucleated CMs. a-actinin (green), DAPI (Blue). Scale bar: 50 μm.

Taken together, we have developed a simple label-free method for isolating highly purified CMs from mouse neonatal hearts and shown that extracted cells retain their functional properties and are suited for down-stream applications such as cardiac drug development and tissue regeneration studies.

## 3. Conclusion

In this work, we used a spiral microfluidic biochip to isolate and purify neonatal CMs from stromal, endothelial and other cell types in the mouse heart. The device performance was evaluated using freshly isolated post-natal day 2 (P2) hearts from C57BL6 mice.

This method offers key advantages such as label free cell isolation, reduced processing time (the whole process < 2 hr), high throughput (up to 120k cells/min), and high purity cell isolation. More importantly, we demonstrated that separation performance of the device can be improved by changing inlet flow rate, changing the location of the outlets by hole punching, and the use of a cascaded setup by simply connecting and running two replicate devices in parallel. This approach allows users to rapidly optimise or fine-tune the device in their laboratory without need to redesign and make a new master mould. CMs from two different transgenic mouse models were used to validate the method by successful demonstration of sort purity and characterisation of the viability, proliferation, immunofluorescence labelling, and contractile behaviour of cells. The utility of the method is not limited to neonatal mouse hearts, but the principles are compatible with isolation of CMs from embryonic, adult and induced pluripotent stem cells derived CMs. The use of spiral biochips is a fast, reliable and relatively cheap tool for isolation and enrichment of CMs. In this study, we have demonstrated for the first-time, the physico-mechanical properties of distinct CMs and mono-nucleated stromal cells in the heart. The mechanical profile of CMs obtained using this method will be useful for tissue engineering purposes where the design of tissue scaffolds with appropriate stiffness to match that of cardiac tissue is necessary.

## 4. Experimental Section

### 4.1. Device design and fabrication

The spiral biochip was fabricated according to [38, 46, 62] has 8-loops, a trapezoidal cross-section with a 600 μm width and the inner and outer heights of 80 and 130 μm, containing one inlet and two outlets. The master mould was designed according to [87] via SolidWorks software and fabricated on an aluminium sheet using the conventional micro-milling method. The mould fabricated by micro-milling process that does not require silanization, it is cost effective and can be reused for extended period. Subsequently, Polydimethylsiloxane (PDMS) pre-polymer and the curing agent (Sylgard^®^ 184, Dow Corning, USA) were mixed, poured into the aluminium mould, degassed in a vacuum chamber for 15 min, and baked at 60 °C for 2 h. Finally, PDMS was peeled off from the mould, inlets and outlets of 1.5 mm diameter were punched with Harris Uni-Core (Ted Pella, Inc., USA), PDMS layers with microchannel features were bonded together after plasma activation.

### 4.2. Neonatal cardiac cell isolation

Neonatal hearts were dissected from wild-type C57/BL6 and *Gt(ROSA)26Sor^tm14(CAG-tdTomato)Hze^* (tdTomato) [88] transgenic mice at postnatal day 2 (P2), and were dissociated into single cell suspensions using a previously published method for adult rat heart cells with some modifications [89]. Briefly, the tissue was minced and digested using digestion buffer containing (mmol/L): 130 NaCl, 5 KCl, 0.5 NaH2PO4, 10 HEPES, 10 Glucose, 10 BDM, 10 Taurine, and 1 MgCl2 supplemented with 0.5 mg/mL Collagenase IV and 0.05 mg/mL Protease XIV at 37°C. The tissue mixture was gently shaken for 30 min until the hearts were completely dissociated. Cardiac cells were isolated by excluding dead cells using the MACS dead cell removal kit and were suspended in 2% FCS in PBS. Fibroblasts exhibit large variation in morphology, size and shapes [90] and are also known to reduce in size during aging [91]. Due to the heterogeneous nature of fibroblasts, fibroblast size may overlap with CM size. Therefore, we used a differential attachment method modified from Yang et al., [12] to rapidly remove fibroblasts. In brief, after the tissue digestion step, cell suspensions were plated onto a plastic culture plate for 1 hour to deplete fibroblasts, which preferentially attach to plastic. The supernatant containing cardiac cells was kept at 4°C until microfluidic sorting. The UNSW Animal Ethics Committee approved all animal experiments.

### 4.3. Device setup

Suspended cardiac cells in 2% FCS in PBS with a concentration of 50,000 cells/ml were loaded into syringes (Terumo Syringes Luer Lock Tip, SSS Australia) and pumped into the spiral biochip using a syringe pump (NE-1800, New Era Pump Systems, Inc., USA) via silicone tubing (Masterflex, Cole-Parmer, USA). Before running the experiment with cardiac cells, the device was washed by Milli-Q water for 5 min to remove any residue and followed by 2 mM EDTA for 5 min to avoid cells sticking to the channel walls. Sorted cells collected from outlets were analysed using the methods described in the next section.

The performance of the spiral biochip was further optimised by adjusting the flow split ratio. The flow split ratio (SP) is defined as follows:

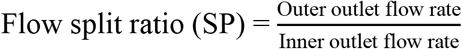

Three spiral biochips with different flow split ratio were designed and denoted as SP1, SP2 and SP3. The master mould was designed with extra-long branching tubes as shown in Figure 3, the same mould was used for microfabrication of all three devices with different flow split ratio. The flow split ratio was modulated by changing the outlet tube length thus changing fluidic resistance. This was carried out by selecting different locations along the branching tubes to punch outlet holes as illustrated in Figure 3. The dimensions of SP1, SP2 and SP3 were listed in Table S1 (supporting information). The length of the inner or outer outlet is measured from the starting point of the bifurcation to the end point of the outlet. The inlet flow rate was maintained at Q = 1.2 ml/min for all the flow split ratio being tested. The flow split ratio of each device was determined by measuring the flow rate of fluid coming out from the inner and outer outlets of the device. Flow split ratios of SP1, SP2 and SP3 were determined to be 2.4, 2.0 and 1.1, respectively.

### 4.4. Cell size analysis and real time deformability cytometry (RT-DC)

Details of the RT-DC setup (AcCellerator, Zellmechanik Dresden, Germany) has been described in detail elsewhere [52–54]. In brief, sorted cells were pumped into a PDMS chip along with a solution of sheath fluid (CellCarrier, Zellmechanik Dresden, Germany). The PDMS chip consists of a 300 μm long channel with a 30×30 μm^2^ squared cross section measurement channel. The cross-section is slightly bigger than the cell diameter, thus cells entering the channel experience hydrodynamic shear and normal stress that causes cell deformation. A high-speed camera in the RT-DC setup captures images of single cells that passed through the measurement channel. Cells collected from the inner and outer outlets were analysed using RT-DC, image processing algorithms (ShapeOut, version 0.9.6) and FlowJo v10.6.1 software to quantify the following cell parameters: cell number, minimum length and height, maximum length and height, minimum and maximum aspect ratio, cross-sectional area (A, μm^2^), perimeter (L, μm), circularity 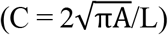, deformation (D = 1-C) [58], as well as Young Modulus.

The separation performance of different devices was compared using percentage of recovery of large cells as an indicator. The percentage of recovery of large cells is calculated as:

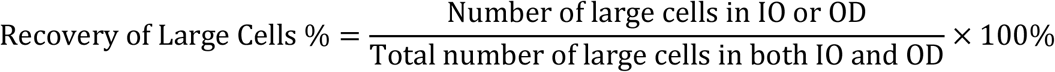

### 4.5. Immunofluorescence staining, and quantification

The cells collected from different outlets were cultured in separated matrigel coated culture plates using culture medium that contained Dulbecco’s Modified Eagle’s Medium (DME)/Ham’s Nutrient Mixture F-12, supplemented with 10 % foetal calf serum (FCS) and Penicillin/Streptomycin (P/S) (10,000 U/mL) in a 5% CO_2_ humidified incubator. Immunostaining was performed to assess the phenotype of cultured cells. After three days of culture, the cultured cells were washed with PBS for 10 min, and then fixed with 4% PFA in PBS (w/v) for 15 min at room temperature (RT). Next, cells were permeabilised with Tween-20 for 15 min, washed with PBS and then blocked with 10% donkey serum in PBS for 2 h. Subsequently, cells were incubated in primary antibodies (detailed in Key Resource Table) followed by three washes with PBS, and final incubation with secondary antibodies for 1 hr. Imaging was performed using a Zeiss LSM 800 confocal microscope. To quantify the separation efficiency, cells expressing α-sarcomeric actinin were counted from three randomly taken images at 20X magnification, and the result was reported as a percentage of CM and non-CM cells in outlets.

### 4.6. Cardiomyocyte beating behaviour analysis

Beating cardiomyocytes were imaged using a Zeiss Axio Observer X.1 SD & TIRF with a 20x phase objective (0.45 NA). 1000 frames were acquired continuously with a 52 ms frame rate. 12-bit images were acquired with a 1280×1024 pixel array. For cell contraction frequency, we created customised software [92] that used normalised cross-correlation to track the displacement of a user specified region over consecutive frames.

## Author’s Contribution

H.T., P.C. and V.C. designed the study. H.T. performed the experiments including microfluidic device fabrication, cardiac cell isolation, cell sorting experiments, RT-DC measurement, microscopic analysis and CM characterization. P.R. and Y.C.K. performed animal mating. H.T. and C.B. performed RT-DC analysis. H.T. and M.C. performed cardiomyocyte contraction analysis. J.P. advised on refining experiments. H.T., P.C. and V.C. analysed and interpreted the data. H.T. wrote the manuscript, which was revised by P.C, V.C. and J.P. P.C. and V.C. supervised this study. P.C., V.C. and J.P. acquired funding.

## Acknowledgements

H.T. thanks the financial support from Swinburne University of Technology through SUPRA scholarship, and the Department of Pathology, UNSW medicine. Funding for this research was partly through Australia Research Council Discovery Project Grant (DP170103704) and UNSW medicine.

