## Supplemental File for "Label-free Isolation and Single Cell Biophysical Phenotyping Analysis of Primary Cardiomyocytes Using Inertial Microfluidics"

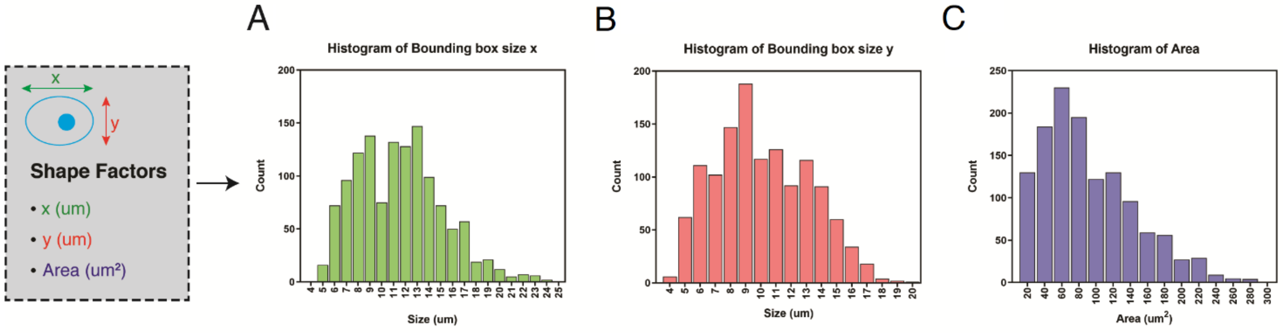


**Figure S1.** Histograms showing the distribution of cell size based on A) longer dimention (x), B) shorter dimention (y), and C) cell area.

### Determination of size threshold for CMs isolation

To find the size threshold for size-based isolation of CMs, we first used a commercially available isolation kit (Miltenyi Biotech, #130-100-825) to isolate neonatal CMs via magnetic activated cell sorting (MACS) method. Neonatal cardiac cells isolate, and non-CM isolate were prepared according to the manufacturer protocol. Next, we measure the size of CMs and non-CMs cells using two methods: i) both populations (CMs vs. non-CMs) were imaged using optical microscope. Their diameter and cross-section area (cell area) were measured using ImageJ software as illustrated in Figure S2 E. ii) CMs and non-CMs populations were separately measured by RT-DC, their dimensions and cell area were determined as illustrated in Figure S2 F. As cells are not identical in shape and are not always spherical in shape, using cell diameter to compare cell sizes will not be appropriate. We therefore used a parameter cell area (A) to compare sizes between different cells.

According to the box and whisker plots and dot plots presented in Figure S2-B and Figure S2-D, CMs have a median cell area of ~125 $\mu m^2$, while non-CM cells have a median cell area of ~ 75 $\mu m^2$. The majority of dots representing CMs laid within the box extending from the 25th to 75th percentiles, whereas more than 75% of the dots representing non-CM cells have a cell area < 100 $\mu m^2$. Based on these observations we selected A=100 $\mu m^2$ as a threshold for size-based isolation of CMs.

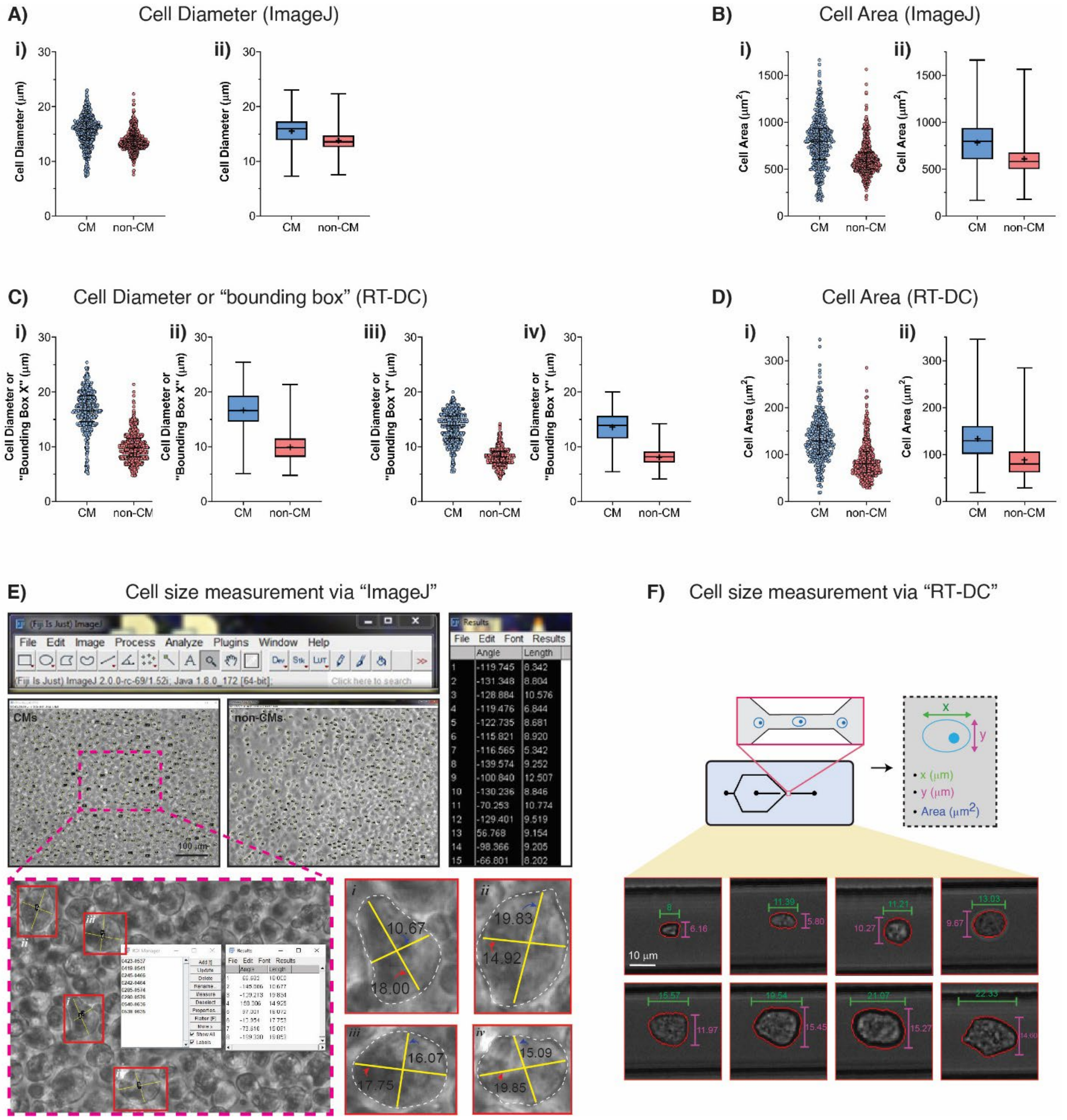


**Figure S2.** Size of CMs and non-CMs. Cardiac cells were harvested from neonatal wild-type C57/BL6 mouse. CMs and non-CMs were isolated using using MACS® method, their sizes were evaluated as shown in **(A-D)** dot plots and box and whisker plots. **(A)** Cell diameter of CMs and non-CMs measured using microscopy imaging and image processing by ImageJ. **(B)** Cell area of CMs and non-CMs as measured using microscopy imaging and image processing by ImageJ. **(C)** Cell diameter or size of cell bounding box. X axis (i, ii) and Y axis (iii, iv) of the cell bounding box as measured by RT-DC. (D) Cell area of CMs and non-CMs measured using RT-DC. The majority of CMs were found to be larger than 100 μm$^2$. All box and whisker plots denote the median within a box extending from the 25th to 75th percentiles and error bars span minimum to maximum values within the indicated datasets. Images in **(E)** and **(F)** illustrate how the cell diameter and cell area were measured using ImageJ and RT-DC, respectively.

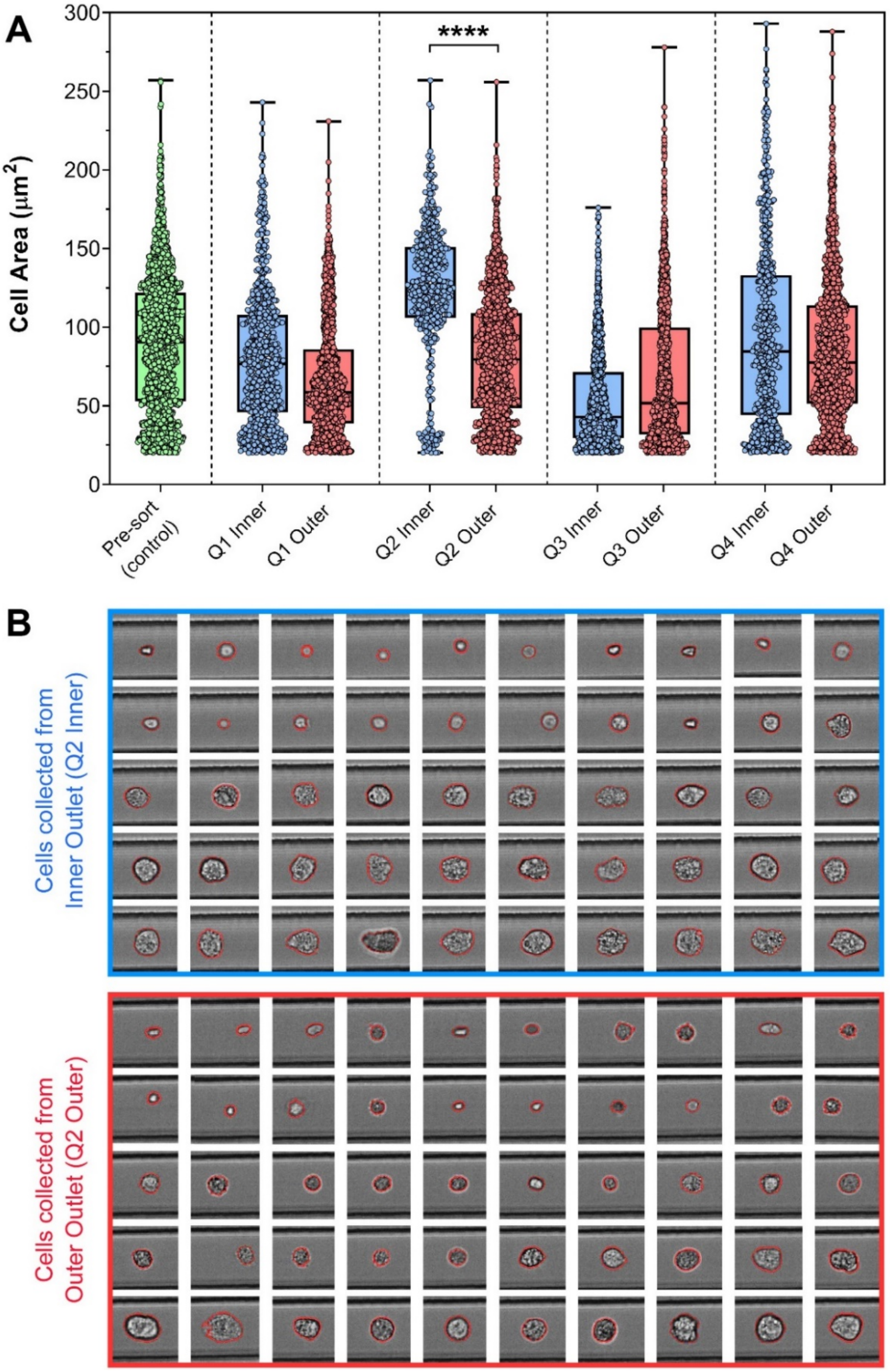


**Figure S3. A)** Box and whisker dot plot showing the size distribution of cells in inlet feed (Pre-sort control, green color) and size distribution of cells in inner outlet (blue color) and outer outlet (red color) after sorting. Four different inlet flow rates were examined: Q1=1 ml/min, Q2=1.2 ml/min, Q3=1.4 ml/min, and Q4=1.6 ml/min. There is a significant difference in size between cells in the inner and outer outlets at Q=1.2 ml/min. The top and the bottom of the box plot represent 75$^{th}$ and 25$^{th}$ percentile values, respectively. The whiskers outside the box represent 10$^{th}$ to 90$^{th}$ percentile values. Data was analysed using two-tailed student's t-test. **** represents $p<0.0001$. **B)** Microscopic images of sorted cells collected from the inner outlet (blue outline) and the outer outlet (red outline) at Q2=1.2 ml/min. The images were taken using RT-DC. All images are at the same scale. Scale bar represents 10 μm.

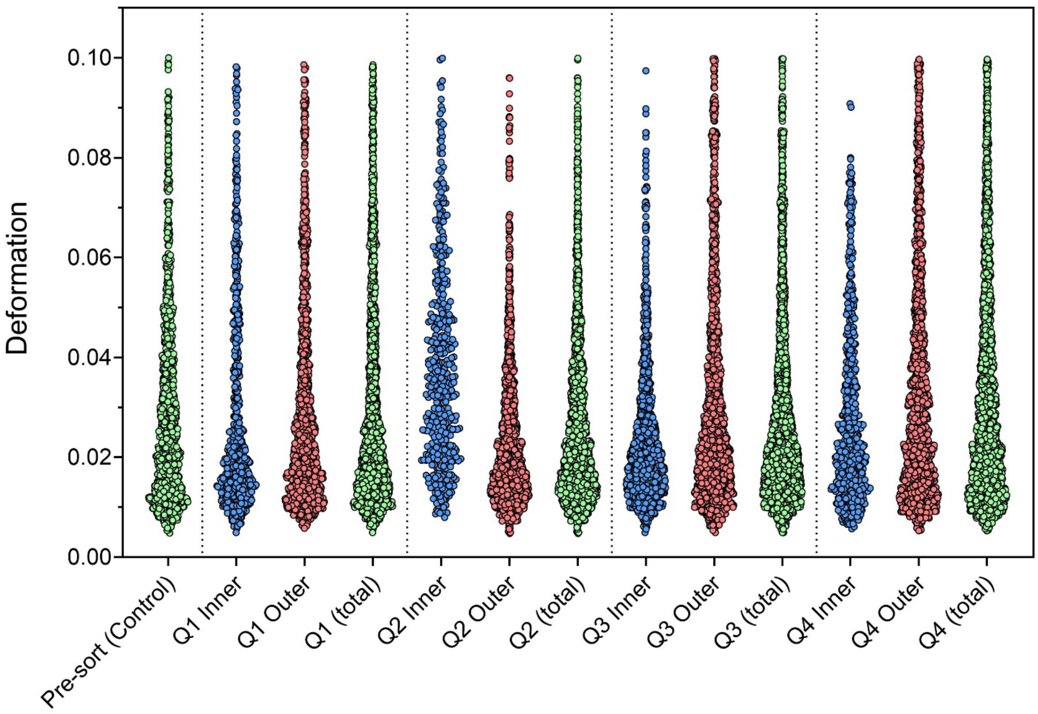


**Figure S4.** Violin plot shows the deformation values of cells before sorting (green dots, control), cells collected from inner outlets (blue dots), cells collected from outer outlets (red dots), and total cell population collected from both inner and outer outlets (green dots). The deformation values were measured at different flow rates: Q1 = 1.0 ml/min, Q2 = 1.2 ml/min, Q3 = 1.4 ml/min, Q4 = 1.6 ml/min.

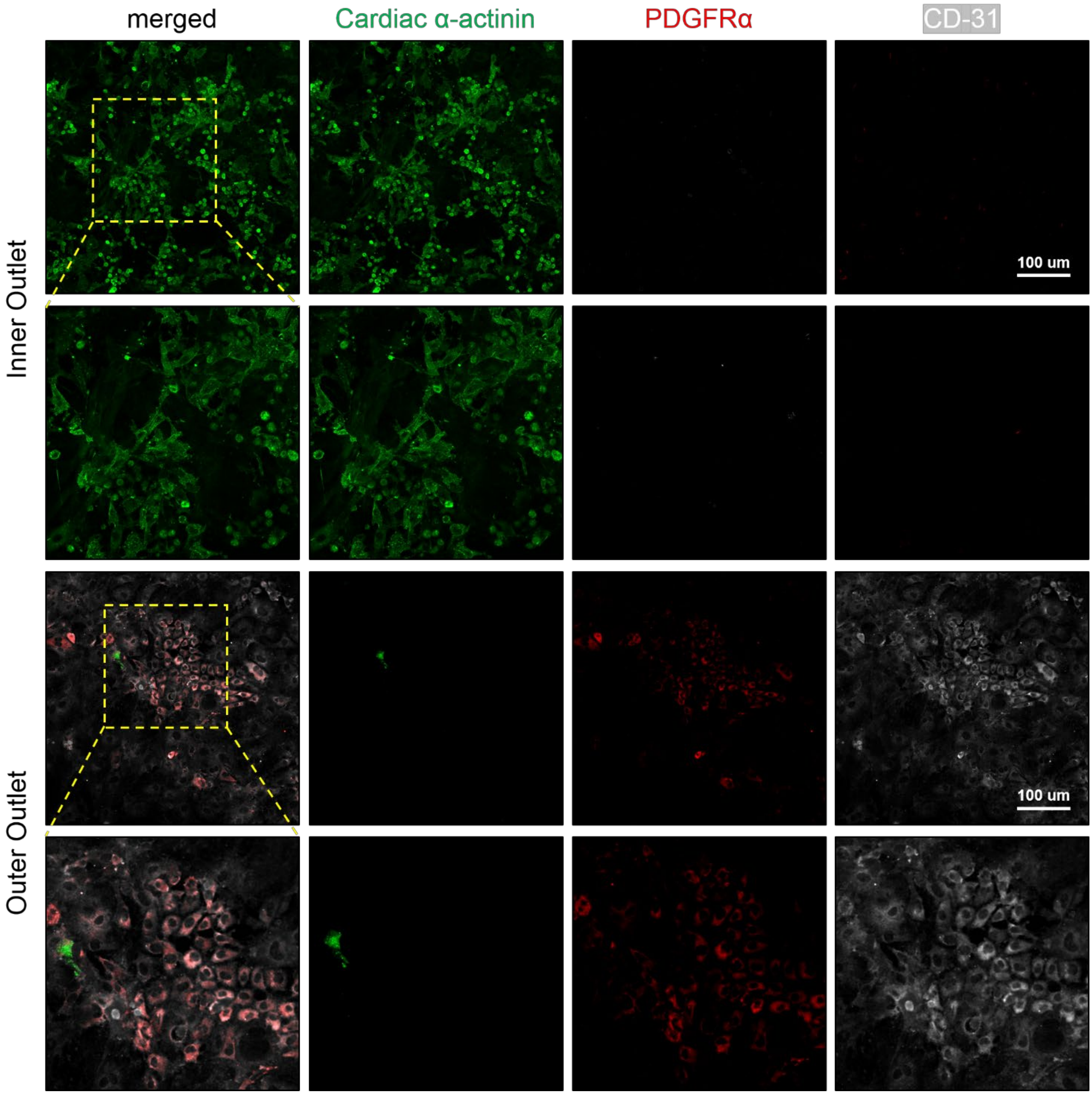


**Figure S5.** Immunofloresnce labelling of sorted nenonatal cardiac cells isolated from C57/BL6 mice into inner and outer outlets. Row 1 and 3 are taken using 10x magnification, and Row 2 and 4 are taken via the 20x.

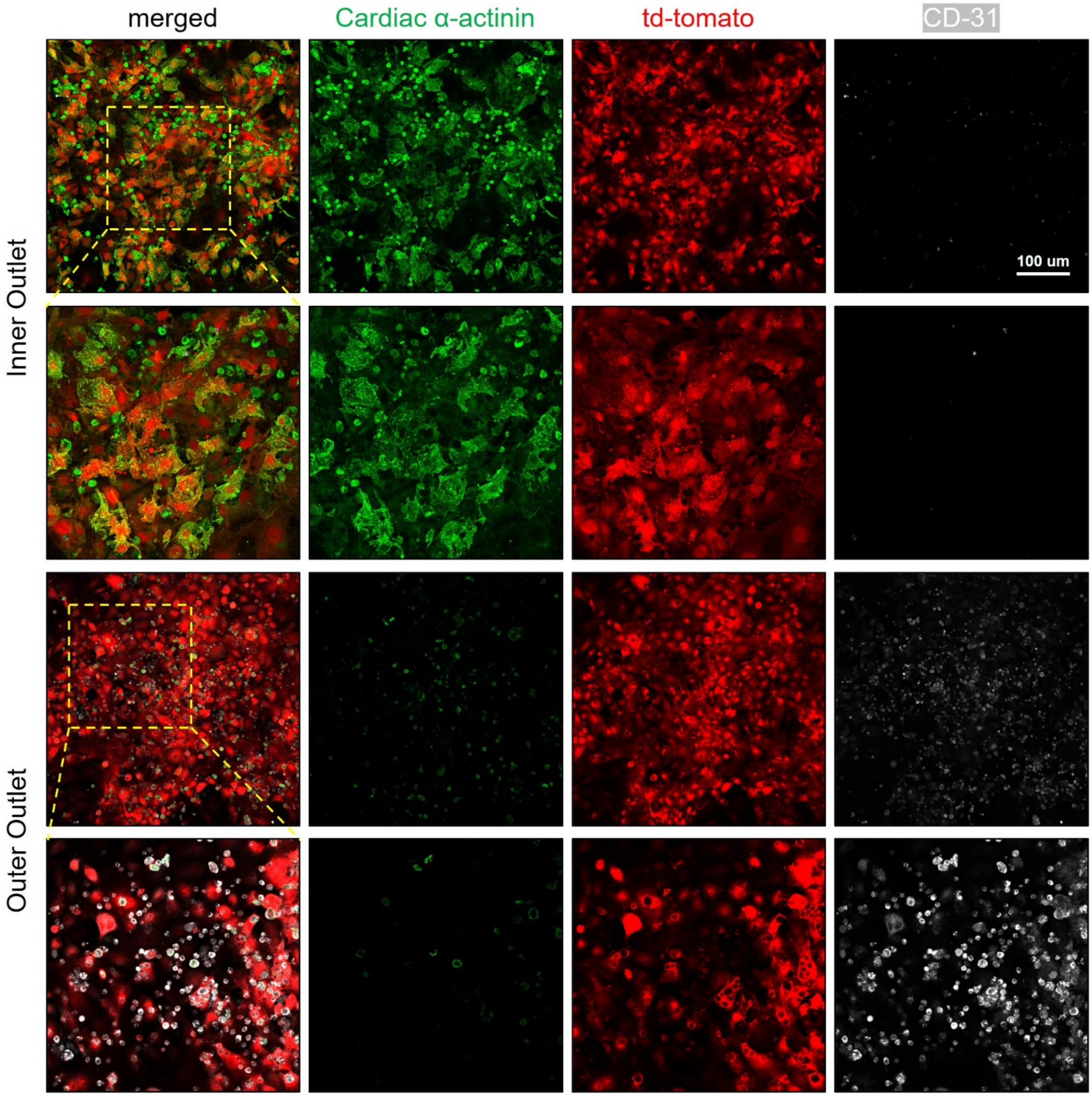


**Figure S6.** Immunofloresnce labelling of sorted nenonatal cardiac cells isolated from transgenic tdTomato mice into inner and outer outlets. Row 1 and 3 are taken using 10x magnification, and Row 2 and 4 are taken via the 20x.

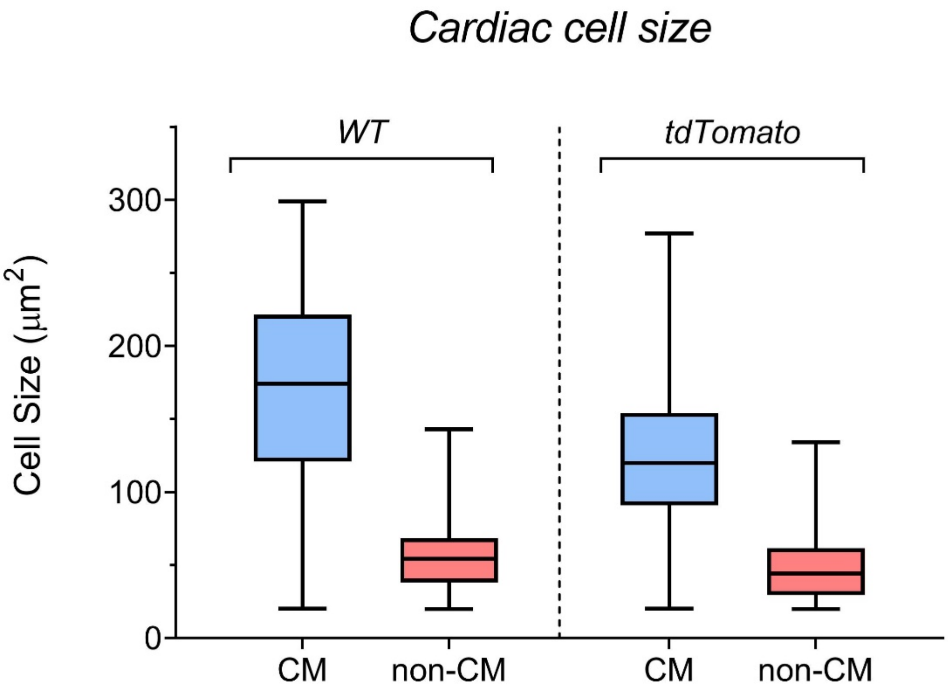


**Figure S7.** Comparison of neonatal cardiac cell size and non-CM cell size in C57BL/6 wild type (WT) mouse and *Gt(ROSA)26Sor*$^{tm14(CAG\text{-}tdTomato)Hze}$ (tdTomato) transgenic mouse. There was a a significant difference in cell size between CMs and non-CMs within the same mice strain. There was also a significant difference in CM cells size between the WT and tdTomato mice. The box and whisker plots denote the median within a box extending from the 25th to 75th percentiles and error bars span minimum to maximum values within the indicated datasets.

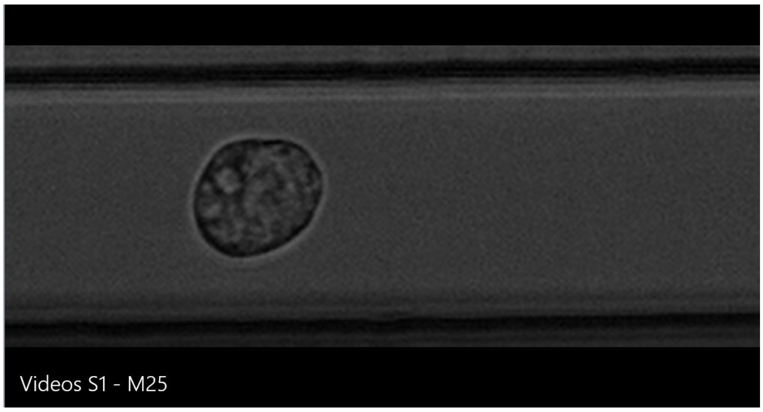


**Video S1.** Analysis of pre-sorted cardiac cells via Real-time deformability cytometry (RT-DC) setup. Scale bar=10 μm.

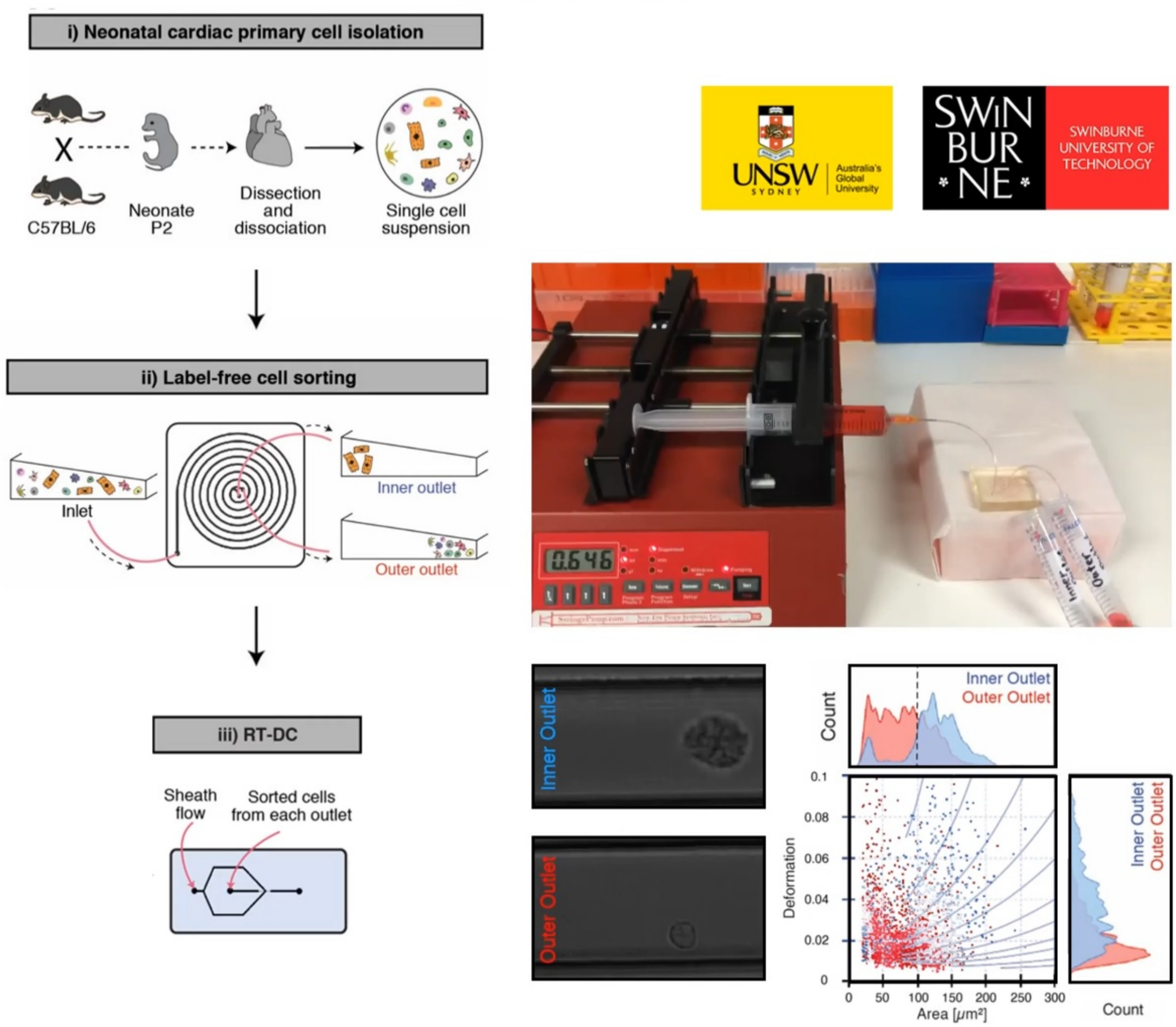


**Video S2.** Cardiac cell sorting using spiral microfluidic biochip at an optimised flow rate of Q =1.2 ml/min, and the cells from the inner and outer outlets were analysed using RT-DC.

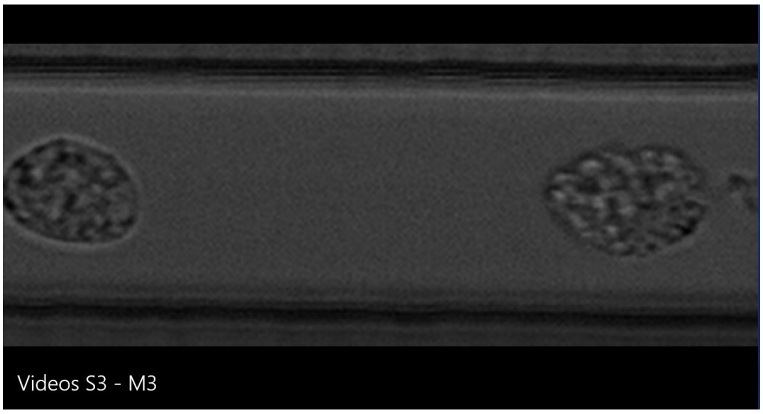


**Video S3.** RT-DC analysis of sorted cells collected from inner outlet of spiral sorting biochip. Cells are mainly large cells i.e., CMs. Scale bar=10 µm.

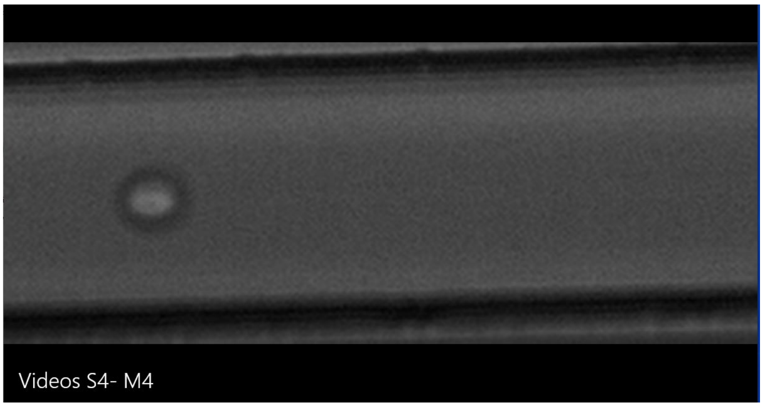


**Video S4.** RT-DC analysis of sorted cells collected from outer outlet of spiral sorting biochip. Cells are mainly small cells i.e., non-CMs. Scale bar=10 µm.

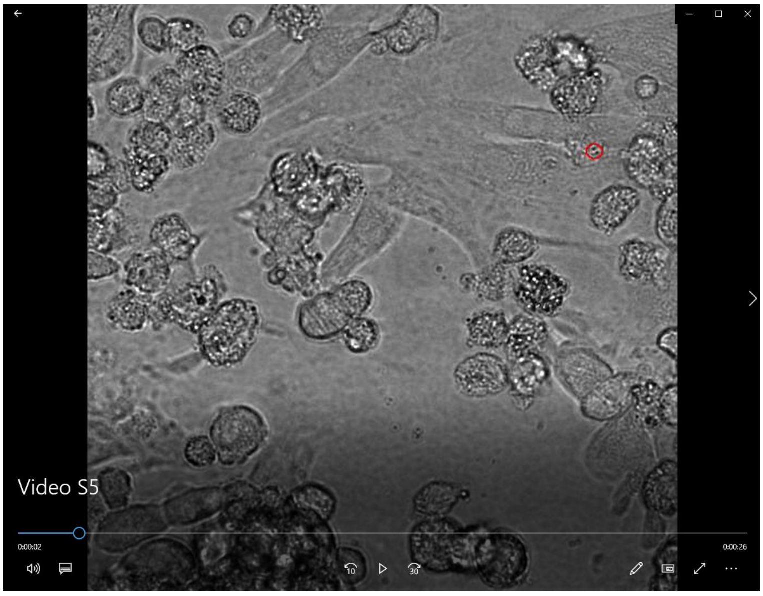


**Video S5.** Live cell imaging showing contraction pattern of CMs (isolated from c57BL/6 mice) after 48 hrs of culture. Scale bars = 20 µm.

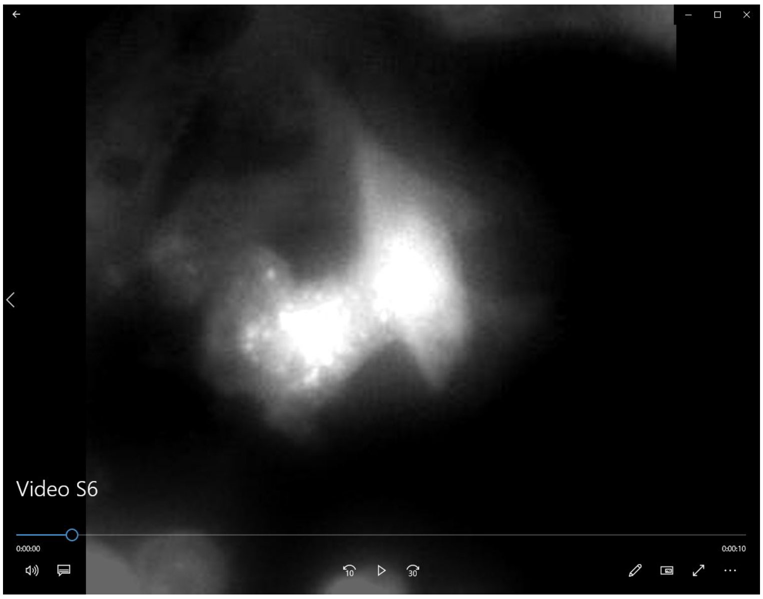


**Video S6.** Live cell imaging showing contraction pattern of CMs (isolated from tdTomato mice) after 48 hrs of culture. Scale bars = 20 µm.

**Table S1.** The ratio of fluid collected from the inner and outer outlet at different split ratio with inlet flow rate Q=1.2 ml/min

| | **Length of inner outlet tube (mm)** | **Length of outer outlet tube (mm)** | **Distance between inner outlet and outer outlet (mm)** | **Volume percent of fluid from inner outlet (%)** | **Volume percent of fluid from outer outlet (%)** | **Split ratio** |
|---|---|---|---|---|---|---|
| SP 1 | 10 | 10 | 5 | 29.4 | 70.6 | 2.4 |
| SP 2 | 5 | 10 | 3 | 33.3 | 66.7 | 2.0 |
| SP 3 | 10 | 5 | 7 | 46.9 | 53.1 | 1.1 |

**Table S2.** Key Resource Table

| REAGENT or RESOURCE | SOURCE | IDENTIFIER |
|---|---|---|
| **Antibodies** | | |
| α-sarcomeric actinin | Sigma-Aldrich | A7811 |
| Anti-CD31 (Rabbit anti mouse) | Abcam | ab28364 |
| Anti-Pdgfra (Rat anti mouse) | BioLegend | 135902 |
| DAPI | H3569 | Invitrogen |
| Donkey anti Mouse-488 | Invitrogen | A21202 |
| Donkey anti Rat-488 | Life Tech | A21208 |
| Donkey anti Rat-650 | Invitrogen | A10043 |
| Chicken anti Mouse-647 | Invitrogen | A21463 |
| Rabbit anti Mouse-555 | Invitrogen | A21427 |
| Donkey anti Rabbit-555 | Life Tech | A31572 |
| **Chemicals** | | |
| Dulbecco's Modified Eagle's Medium (DME)/Ham's Nutrient Mixture F-12 | Sigma-Aldrich | D6421 |
| Penicillin/Streptomycin (P/S) (10,000 U/mL) | Thermofisher Scientific | 10378-016 |
| Fetal Calf Serum (FCS) | Scientifix | NZFBS-25 |
| Bovine Serum Albumin (BSA) | Sigma Aldrich | A2153 |
| Donkey serum | Sigma Aldrich | D9663 |
| MACS dead cell removal kit | Miltenyi Biotec | 130-090-101 |
| Matrigel Matrix | Corning | 356237 |
| Paraformaldehyde (16%) | Electron Microscopy Sciences | 15710 |
| **Software and Algorithms** | | |
| FlowJo | Tree Star | v10.6.1 |
| GraphPad Prism | GraphPad Software, Inc. | v7 |
| ShapeOut | Zellmechanik Dresden | 0.9.6 |
